# A generalizable codesigned platform for solid-state nanopore sensing beyond the capacitive-noise constraints

**DOI:** 10.64898/2026.07.06.731876

**Authors:** Nianduo Cai, Wei Guo, Yunfei Teng, Yameng Lou, Siu-Ho Wong, Akhil Sai Naidu, Filippo Cona, Federico Thei, Tzu-Heng Chen, Maartje Bastings, Aleksandra Radenovic

## Abstract

Solid-state nanopores offer label-free, real-time single-molecule sensing, yet resolving rapidly translocating biomolecules requires high-bandwidth data acquisition, where increased high-frequency noise fundamentally limits reliable recovery of informative events. Here we establish a hardware-software co-designed nanopore sensing platform that overcomes this bandwidth-noise limitation by integrating scalable low-noise device engineering with deep learning-based signal reconstruction. A wafer-scale dielectric-engineering strategy using low-dielectric SU8 photoresist reduces total device capacitance to the picofarad regime and suppresses high-frequency noise by up to fivefold while maintaining facile, controllable and reproducible fabrication. This extends usable acquisition rate to 40 MHz and enables capture of fast molecular features. Combined with a reconstruction neural network trained on synthetic translocation signals embedded in experimentally measured noise, the platform recovers transient blockade sublevels while preserving temporal fidelity. Using engineered DNA molecules carrying dumbbell-like barcodes, we resolve nanometer-scale structural features on sub-microsecond timescales, and experimentally measure translocation velocity within the sub-5 nm regime. Dual-channel measurement on a single nanopore device further demonstrates transferability of the platform by showing robust cross-channel signal reconstruction across distinct baseline noise levels. This platform provides a general route for reliable recovery of previously inaccessible molecular information from high-bandwidth nanopore measurements.

## Main

Nanopore sensing has emerged as a single-molecule analysis platform that enables label-free, real-time electrical detection of biomolecules^1–4^ and engineered molecular constructs^5–8^. As nanopore sensors essentially converts minute structural and chemical information of the analytes into ionic current signals, the ability to decode transient event features with high precision becomes a fundamental determinant of its utility. This remains challenging because many informative nanopore events occur on timescales exceeding conventional measurement bandwidth (typically from hundreds of kHz to a few MHz) due to the fast electrophoretic translocation through the pore^4,9–11^, causing short-lived events and closely spaced signal features to merge or become unresolved. Various strategies have therefore been developed to prolong the event duration to relax the bandwidth requirements, including enzyme-assisted control of nucleic-acid transport in biological nanopores^12–15^ and modulation of pore geometry^16–19^ or electrolyte conditions^10,20,21^ in solid-state nanopores. Yet these methods improve signal observability primarily by altering molecular transport dynamics rather than preserving the intrinsically fast event features that naturally encode molecular information.

Solid-state nanopores remain particularly attractive as they offer high stability, compatibility with scalable microfabrication and flexibility in device engineering, making them promising candidates for next-generation single-molecule sensing technologies^22–24^. Nevertheless, their practical performance is fundamentally constrained by the trade-off between temporal resolution and electrical noise. Capturing fast translocation features requires high-bandwidth acquisition, while increasing bandwidth amplifies high-frequency noise and obscures the fine structural details embedded in the signal^25–27^. This bandwidth-noise trade-off becomes significant when decoding engineered molecular barcodes, where reliable readout depends on resolving rapid and closely spaced features programmed on DNA carriers^28,29^.

Considerable efforts have therefore focused on improving nanopore signal fidelity through both device engineering and computational analysis. On the hardware side, reducing chip capacitance and suppressing parasitic contributions have been applied to lower high-frequency noise^25,27,30,31^. In parallel, machine learning and deep learning-based approaches have shown increasing promise for signal denoising^32,33^ and event classification^34–38^ from complex nanopore recordings. Despite these important advances, a scalable and transferable end-to-end platform that integrates low-noise, high-bandwidth acquisition with robust computational recovery of fast molecular features across varying experimental conditions remains an unmet need.

Here, we develop such a hardware-software co-designed platform for reliable recovery of transient features in high-bandwidth nanopore signals (**Figure 1a**). We first develop a wafer-scale, dielectric-engineered solid-state nanopore sensor that significantly suppresses high-frequency noise. To achieve this, we establish a facile and scalable microfabrication strategy that enables reproducible integration of low-dielectric constant SU8 photoresist on the silicon nitride (SiN_x_) membrane of nanopore chips. This dielectric engineering reduces the total capacitance of the measurement system to a few picofarads and suppresses high-frequency noise by up to 5-fold compared with conventional SiN_x_ nanopore. The controllable fabrication method further enables systematic investigation of the interdependence between dielectric architecture, device capacitance and electrical noise characteristics. Together, these improvements extend the measurement bandwidth to 10 MHz and enable direct recording of molecular translocation signals at sampling rates up to 40 MHz, providing sufficient temporal resolution to capture short-lived signal features corresponding to engineered DNA barcodes. Building on this low-noise high-bandwidth data acquisition capability, we then introduce a deep learning (DL)-based signal reconstruction framework trained on experimentally measured noise, to fully exploit the enhanced temporal bandwidth. The combined hardware-software platform enables reliable recovery of barcode features spaced within a few nanometers on free-translocating double-stranded DNA backbones with high-speed recordings.

**Figure 1.**
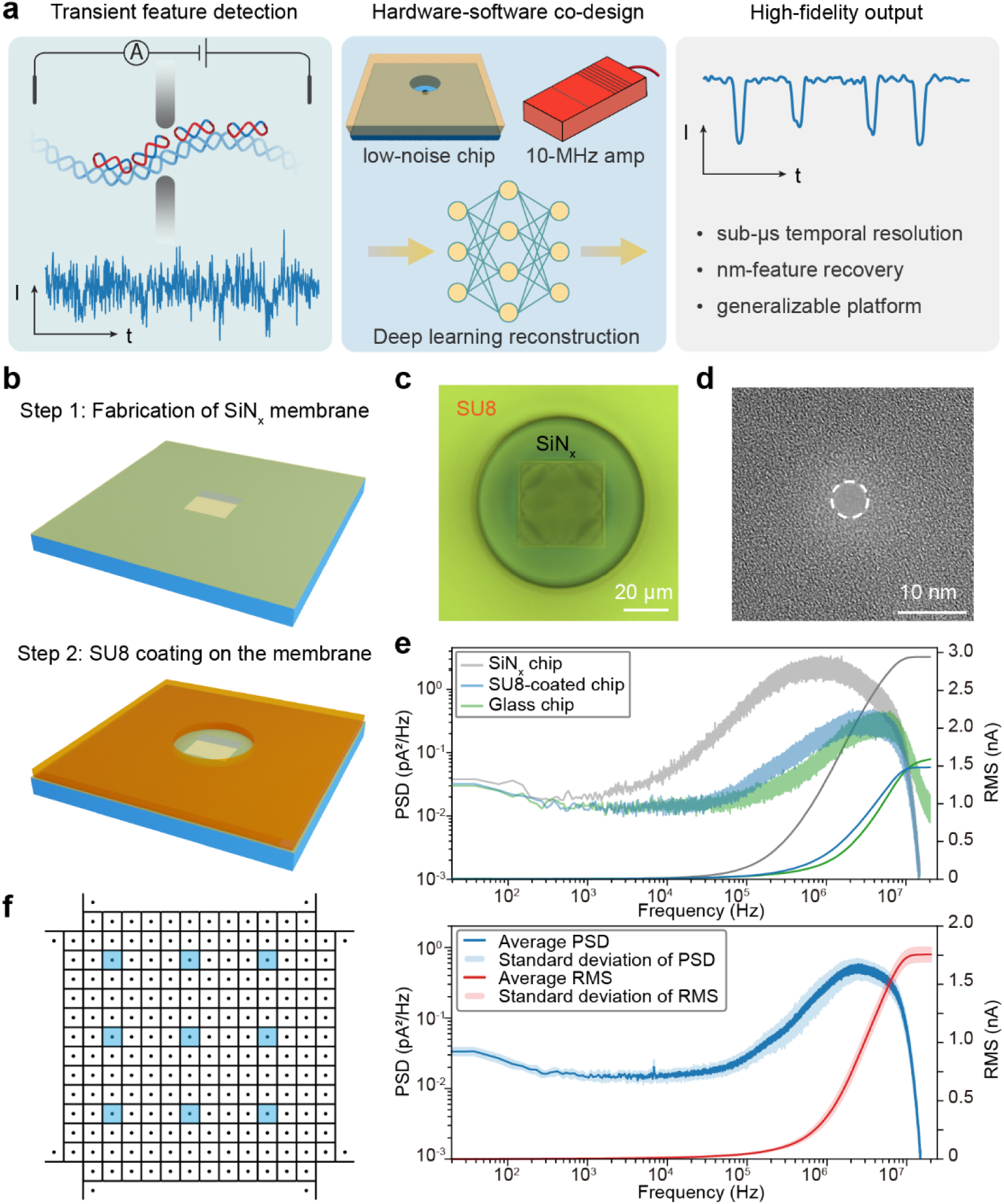
High-bandwidth sensing enabled by low-noise solid-state nanopore device. **(a)** The detection of transient features on molecules is enabled by a co-designed platform, including low-noise nanopore chip and high-bandwidth amplifier on the hardware side, and a deep learning-based reconstruction model on the software side. **(b)** Schematic overview of the fabrication process at wafer-scale. From step 1: silicon chips (in blue) are prepared with a suspending window in the centre of 20 nm-thin silicon nitride (SiN_x_) membrane (in yellow). Step 2: A layer of SU8 is coated on the top surface, and an aperture with tunable size is fabricated in the coating for liquid access to the SiN_x_ membrane. **(c)** Optical image of an SU8-coated chip. Two images with different focal lengths are stacked to have a clear view of SiN_x_ membrane and top rim of SU8 aperture due to the high aspect ratio of SU8 aperture. **(d)** TEM image of a nanopore with 4-nm in diameter in SiN_x_ membrane. **(e)** Baseline current power spectral density (PSD) and root-mean-square (RMS) noise of bare SiN_x_ chip, SU8-coated chip and commercial glass-substrate chip. Here, the SU8 aperture size is 40 µm in diameter and thickness is 40 µm. **(f)** Wafer-scale stability of noise reduction performance. 9 devices (in blue) from different locations on the wafer (Left) were tested, with mean and standard deviation of PSD and RMS noise computed (Right). The SU8 aperture size here is 80 µm in diameter and thickness is 40 µm. All the noise characterization measurements are performed in 1 M KCl at 10 MHz bandwidth and 40 MHz sampling rate via Elements instrument, with applied voltage bias of 100 mV.

We further demonstrate the transferability of this platform by simultaneously recording the same molecular translocation events on two independent channels with distinct baseline noise characteristics, and reconstructing both signals with the same deep-learning framework. The consistent recovery of fine molecular features across matched events shows that the method is robust to different experimental noise conditions. Altogether, these results establish a generalizable strategy for high-bandwidth nanopore sensing beyond the bandwidth-noise limitation and extend the resolvable information content from single-molecule measurements.

## Results

### Wafer-scale noise reduction

To fabricate the low-noise nanopore devices (**Figure 1b**), we start with a silicon chip that presents a suspended 40 µm x 40 µm window in the central region of the 20 nm-thin SiN_x_ membrane (Step 1, detailed protocols in **Supplementary Fig.1**). A continuous layer of SU8 photoresist is then spin-coated on the membrane and patterned by a lithography process to leave a central aperture on the membrane (Step 2). The SU8 aperture aligns vertically with the SiN_x_ membrane to enable liquid access to the nanopore (**Figure 1c**). Since both steps are processed at wafer-scale, 192 chips are produced in one batch and higher yield can be realized with refined lithographical design (**Supplementary Fig.1**). Moreover, leveraging the versatility and compatibility of SU8 in lithography-based fabrication method, we additionally fabricate apertures with truncated conical shapes via two-photon greyscale lithography to further suppress high-frequency noise by reducing the electrolyte-exposed area of SiN_x_ membrane (**Supplementary Fig.2**). Nanopores with a range of 5-7 nm in diameter are produced on the suspending SiN_x_ window using Transmission Electron Microscope (TEM) (**Figure 1d**).

The noise level is then characterised in 1 M KCl to facilitate comparison with existing studies as it is most commonly used^39–41^, and we compare three different types of solid-state nanopore devices with a 10 MHz amplifier at 40 MHz sampling frequency, including the bare SiN_x_ chip without coating, SU8-coated chip and commercial glass-substrate chip. The SU8 coating has greatly suppressed the noise power in frequency domain of over 10^3^ Hz (**Figure 1e**), which is a typically defined regime for dielectric noise and capacitive noise^26^. In the same measurement configuration, the SU8-coated chip presents a comparable noise level as the glass-substrate chip, which is a benchmark low-noise solid-state nanopore.

The SU8-coated chip is also compared with other state-of-the-art noise-reduction methods with different instruments and shows over 5-fold noise reduction under 200 kHz sampling frequency, indicating a broad effectiveness from mid-high to high bandwidth acquisition (**Supplementary Fig.3**). Given the complicated fabrication protocol of glass-substrate nanopore and uncontrollable silicone painting method, the SU8 coated chip presents convenience and higher reproducibility. We further test the wafer-scale stability of noise-reduction performance by measuring 9 chips from different locations (**Figure 1f**), and approximately 2% relative deviation in noise level is shown at the highest sampling bandwidth, indicating good reproducibility of noise-reduction across the wafer. The difference could potentially originate from slight deviation in thickness of the coated layer, as shown in the wafer-scale SU8 thickness mapping result in **Supplementary Fig.4**.

### Device noise and capacitance modulation

The dominant noise sources in nanopore system can be separated by frequency, with low-frequency 1/f noise (below ∼100 Hz), intermediate white noise (∼100 – 1000 Hz), and high-frequency dielectric (∼1 – 10 kHz) and capacitive noise (over ∼10 kHz) that becomes significant as bandwidth increases^39,40^ (**Figure 2a**). In this high-frequency regime, chip capacitance and parasitic contributions from the system dominate the measured noise. Previous study showed that nanopore can be modelled as an ionic resistor along with a capacitor with contributions from different layers of materials^31^, therefore introducing an additional SU8 layer with low-dielectric constant should reduce the overall chip capacitance and thus suppress the noise floor.

**Figure 2.**
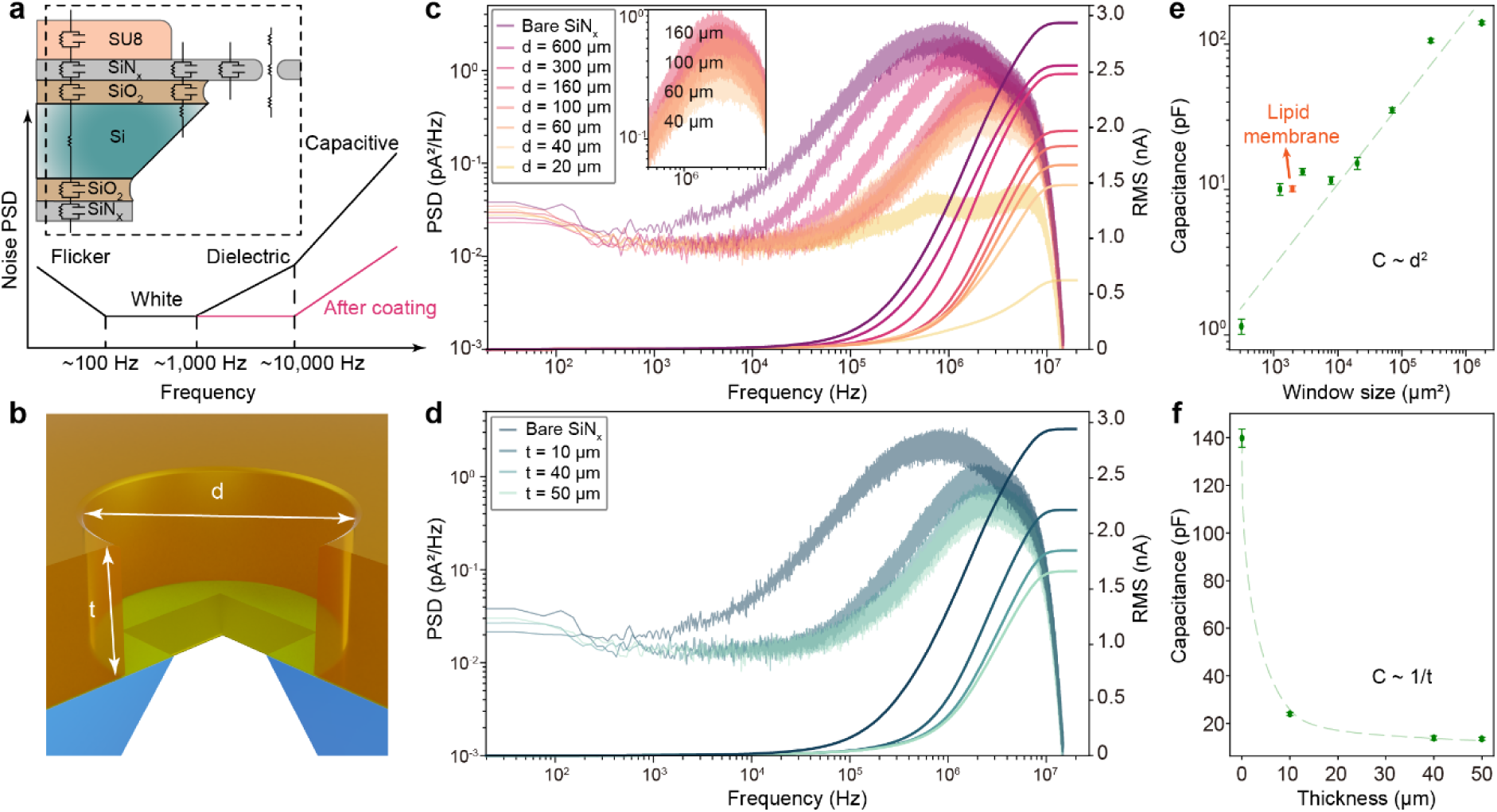
Electrical noise performance and capacitance tuning by SU8 geometry. **(a)** Frequency-dependent noise components illustrated in PSD and cross-section schematic of an SU8-coated chip with equivalent circuit showing the electronic impedances. **(b)** Schematic of the SU8 (yellow) aperture geometry over the SiN_x_ membrane (green) and silicon substrate (blue). The diameter of the SU8 aperture is noted as d and the thickness of coating as t. **(c, d)** PSD and RMS noise as a function of aperture diameter and SU8 thickness. **(e, f)** Measured system capacitance as a function of aperture diameter and SU8 thickness. The dashed lines are visual guides for the trend of capacitance change. The coating thickness is set at 50 µm when tuning the aperture size, while tuning the thickness, aperture size is 80 µm. Noise characterisations are performed in 1 M KCl at 10 MHz bandwidth and 40 MHz sampling rate via Element instrument, with applied voltage bias of 100 mV. The capacitance is extracted from triangle-wave test at 1 MHz bandwidth with voltage amplitude of 100 mV and sweeping frequency of 5 Hz.

The lithography-based fabrication method enables a systematic study to understand experimentally how the SU8-coating reduces noise. By varying the SU8 aperture diameter and the coating thickness (**Figure 2b**), we measure the noise level under the same condition and compute the system capacitance via triangle-wave measurement (**Supplementary Fig.5**). Consistent with the hypothesis, we observe reduction of high-frequency noise starting from the dielectric regime after coating the SU8 layer (**Figure 2c**). As the aperture size becomes smaller, the noise floor is further suppressed as a result of the large area of SU8 layer and smaller area of SiN_x_ membrane in contact with the liquid environment. The reduced exposure of this thin membrane is nontrivial as it contributes to a major part in the chip capacitance (**Supplementary Fig.6**). Alternatively, increasing the SU8 thickness also reduces the high-frequency noise as it dominates the total capacitance of the chip (**Figure 2d**). The measured capacitance from triangle-wave test follows the same trend and indicates how it varies with SU8 geometry (**Figure 2e,f**), in agreement with the equivalent electronic circuit model. It is worth noting that further increasing the thickness could reduce the capacitance while the decreasing trend becomes slow as the limiting factor is on the silicon nitride membrane, and high aspect ratio of SU8 aperture could also lead to practical working problems such as nanopore device wetting. We also calculated the chip capacitance corresponding to each geometry based on the equivalent electrical model, and the results exhibit the same trend of change when the SU8 aperture size and thickness are varied (**Supplementary Fig.6**). These results provide for the first time experimental evidence of how a low-dielectric constant coating layer could controllably modulate the noise in measurements and establish a practical design rule for tuning the capacitance via controlling exposed membrane area and dielectric coating.

### Multi-scale denoising network for feature-preserving signal reconstruction

The SU8 coating has substantially improved noise performance and extends the usable bandwidth of solid-state nanopore readout, enabling the capture of the transient features on the fast translocating molecules. However, the measured traces under high sampling rate still contain residual high-frequency fluctuations that can obscure transient sublevels, making it less certain to correlate detected signals with physical structural features that are closely spaced. In addition, the intrinsic noise floor of 10 MHz-bandwidth data acquisition results in a low signal-to-noise ratio even with noise-reduced nanopores (**Supplementary Fig.7**). Conventional digital-processing methods such as down-sampling and low-pass (LP) filtering can improve the visual interpretability, while at the cost of temporal resolution and often lead to biased peak calling results that are cutoff frequency dependent. Therefore, recovering the embedded information computationally after high temporal bandwidth acquisition without compromising event fidelity is required.

To address this bottleneck, we developed a multi-scale denoising network that reconstructs event structure while preserving blockade edges and sublevels (training and prediction process in **Supplementary Fig.8**). As experimentally recorded ground truth labels are usually lacking in nanopore experiment, we adapted a supervised denoising network inspired by previous work^42,43^. The model is thus trained on synthetic translocation events with simplified current blockage profiles that are edge-softened to reflect the finite measurement bandwidth and then overlaid with experimentally measured nanopore noise from baseline recordings (**Figure 3a**). To superimpose real-life noise, a PSD-based analysis is used to decompose the noise into four physical sources according to the frequency signature. RMS noise is then estimated by integrating the noise power in each frequency band and added on the simplified blockage profiles. The network is based on a U-Net architecture (**Figure 3b**), with multi-scale filtered event traces as inputs and an edge-preserving loss function is incorporated. This multi-scale filtering-assisted data preparation and network training strategy allows the network to learn both the intrinsic event morphology and noise characteristics of the experimental system, and it also ensures the reconstructed output to retain both event fidelity and boundaries of blockage profile instead of over-smoothing the traces (**Supplementary Fig.9**).

**Figure 3.**
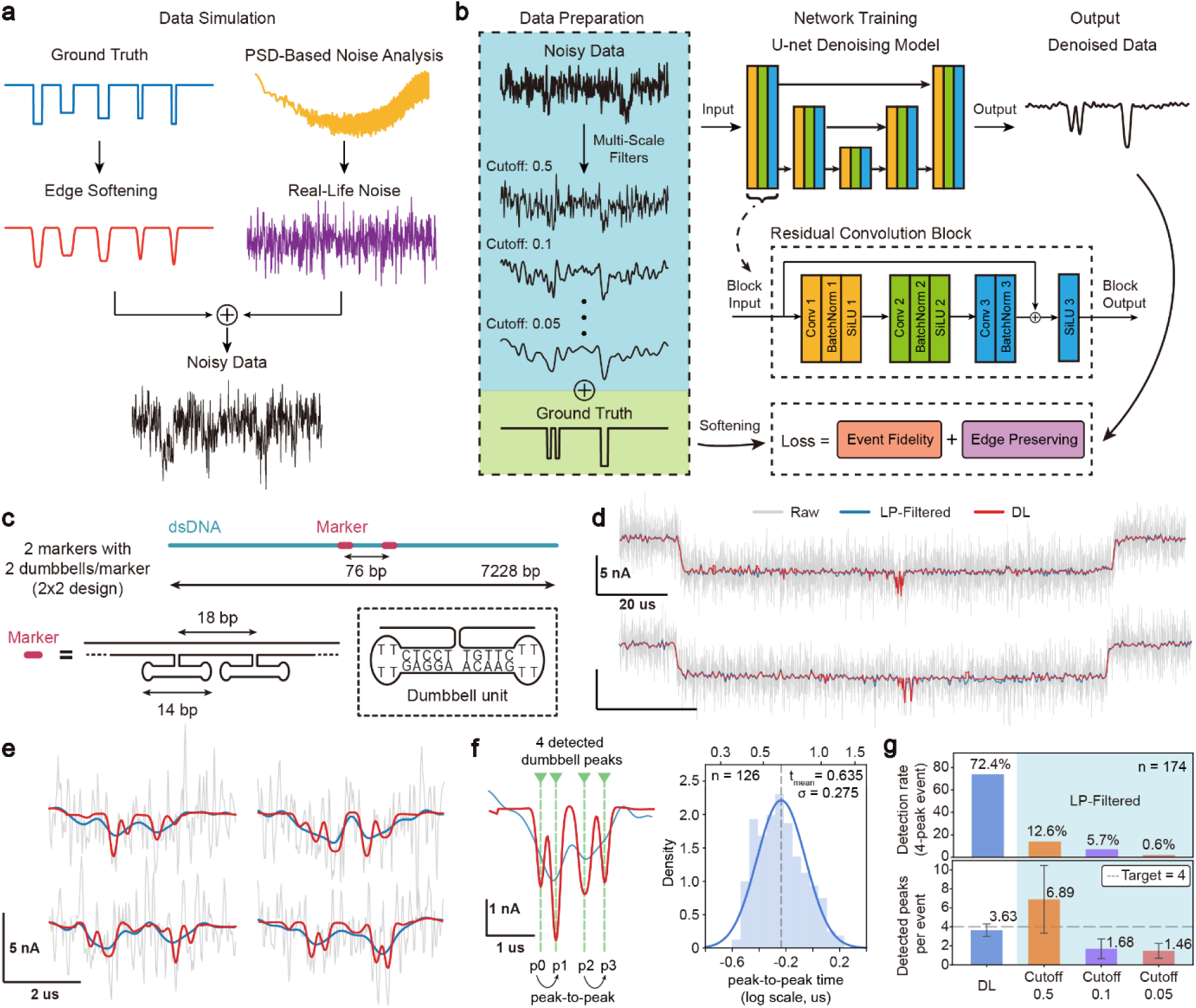
Multi-scale deep-learning reconstruction of fast DNA dumbbell translocations. **(a)** Schematic illustration of synthetic data simulation. **(b)** The multi-scale denoising network architecture and the schematic of its training process. **(c)** Design of DNA molecules with barcode structures. The distances between two markers and the two dumbbells inside each marker are based on the centre-to-centre distance of the structural unit according to the designed DNA sequence. Length of each dumbbell unit represents the full-scale span of the dumbbell side arms. Inset shows the detailed sequence of a single dumbbell unit and the self-paired structure. **(d)** Representative translocation events of the nanostructured DNA molecules in 3 M LiCl solution through a 7 nm-in diameter pore under 1000 mV voltage bias. All three traces are superimposed on same events, including the raw trace (in grey) collected under 10 MHz bandwidth and 40 MHz sampling rate, low-pass (LP)-filtered trace (in blue) with cutoff frequency at 2 MHz, and deep-learning (DL) reconstructed signal (in red). **(e)** Zoomed-in view of translocation signals in marker and dumbbell region, with same colour indication as in **(d)**. **(f)** Example of peak detection result (green dashed line) from the DL reconstructed signal. Each peak is defined with an id according to order. Peak-to-peak time interval is statistically computed using the peak pairs p0-p1 and p2-p3 (252 pairs from n = 126 events) due to the same designed structural gap on the DNA backbone. The peak-to-peak time distribution is plotted in log-scale and fitted with Gaussian (dark blue line). Bottom and top axes are in logarithmic and linear scale, respectively. **(g)** Statistical comparisons of DL reconstruction and LP-filtering in terms of peak calling efficacy (n = 174 events). Three LP-filtering parameters are compared, with 0.5x, 0.1x and 0.01x cutoff frequency. Same peak finding algorithm is applied for peak extraction. Detection rate is calculated on the number of events that have 4 peaks detected among all events. For example, n (4-peak event) = 126 among n (total event) = 174 for the DL-reconstructed result leads to around 72.4% detection rate. Detected peaks per event is estimated and shown with average value and error bar represents standard deviation.

We first benchmarked the model using a nanostructured DNA with closely-spaced dumbbell barcodes (**Figure 3c**). The design is validated in previous study and consists of two marker regions that are separated by a centre-to-centre distance of 76 bp on a double-stranded DNA (dsDNA) backbone with 7228 bp contour length^28^. Each marker was composed of two dumbbell units, with 18 bp distance between the centre of each pair of dumbbells. The dumbbell unit is formed by incorporating pairing nucleotides on barcode sequences which automatically self-assemble into such a hairpin structure. We synthesized the nanostructured DNA molecules via hybridising a single-stranded DNA backbone with the oligonucleotides carrying complementary as well as the hairpin-forming sequences (**Methods** and **Supplementary Information Section 3, Supplementary Table 1,2**) and then measured the free translocation of this DNA construct through SU8-coated low-noise nanopore with pore diameter between 6-7 nm in 3 M LiCl solution at pH 8.0. The ionic current signals were recorded using an amplifier with 10 MHz bandwidth and at 40 MHz sampling frequency, followed by direct DL reconstruction on the raw trace.

**Figure 3d,e** shows the characteristic current blockage signal from the barcoded DNA molecules, in accordance with the reported results on the same molecule translocation measurement using a glass-nanocapillary pore. When applied to the raw 40 MHz signals, the network reconstructs the primary dsDNA backbone blockage together with the secondary dumbbell-associated features that are obscured by residual noise in the raw signal. Moreover, the reconstructed signals match with the LP-filtered reference trace while preserving the sharper sublevel peaks, indicating that the model recovered event features without sacrificing the temporal details carried in high-bandwidth data.

We further analysed quantitatively the dumbbell peak-to-peak time intervals based on the reconstructed traces. The 4 dumbbell-associated secondary peaks are detected first with a peak-finding algorithm (**Methods** and **Supplementary Fig.14**) and labelled from p0 to p3, with p0-p1, p2-p3 representing the two pairs of dumbbells with same dumbbell centre-to-centre distance in each marker. The peak-to-peak time is therefore defined as shown in **Figure 3f** and corresponding time intervals are statistically summarized across n = 126 events, resulting in t (peak-to-peak) = 0.64 ± 0.28 µs after Gaussian fitting on the histogram. If considering the dumbbell centre-to-centre distance as the corresponding molecule travel distance, an average translocation velocity of around 28 bp/µs is obtained, matching the reported typical electrophoretic velocity of nucleic acid across solid-state nanopores (1-100 bp/µs)^1,44^. It is worth noting that, enabled by the extended measurement bandwidth due to reduced device noise and high-bandwidth amplifier, the DL reconstructed signal has reached sub-µs temporal resolution when differentiating two barcodes that are a few -nm separated in space, or a few -bp separated between the side arms of neighbouring dumbbell units. In addition, we also compared the DL-reconstruction results with LP-filtering results under different three filtering levels by computing 4-peak event detection rates and detected peaks per event in each scenario using the same peak-finding algorithm (**Figure 3g**, example events in **Supplementary Fig.15**), the DL-based approach presents more robust peak calling efficacy compared with LP-filtering, and the latter shows cutoff-dependent bias on peak detection.

### Resolving DNA molecules with different barcode density

Having demonstrated the performance of the DL reconstruction on high-bandwidth recordings, we next explored the generality of the same platform by detecting DNA molecules with varying dumbbell units and dumbbell density in individual markers. Two additional constructs with six marker regions are designed (**Supplementary Table 3,4**), with design 1 containing four dumbbells per marker and design 2 containing eight dumbbells per marker. The marker regions across two constructs are identical while the density of dumbbells in the marker varies among the two designs, resulting in 40 bp intra-marker dumbbell centre-to-centre distance in design 1 and 20 bp distance in design 2 (**Figure 4a**). We also modified the single dumbbell unit by adding two C-G pairs to improve the stability of dumbbell hairpin structure **(Supplementary Table 5** and **Supplementary Fig.12,13**). Translocation events of these two molecules were detected with a 7 nm-in diameter low-noise nanopore device in 3 M LiCl solution (pH 8.0) and recorded signals with same amplifier under maximum sampling frequency, followed by DL reconstruction on raw trace using the exact same trained model weights and peak detection method as the previous experiment on the 2-marker sample.

**Figure 4.**
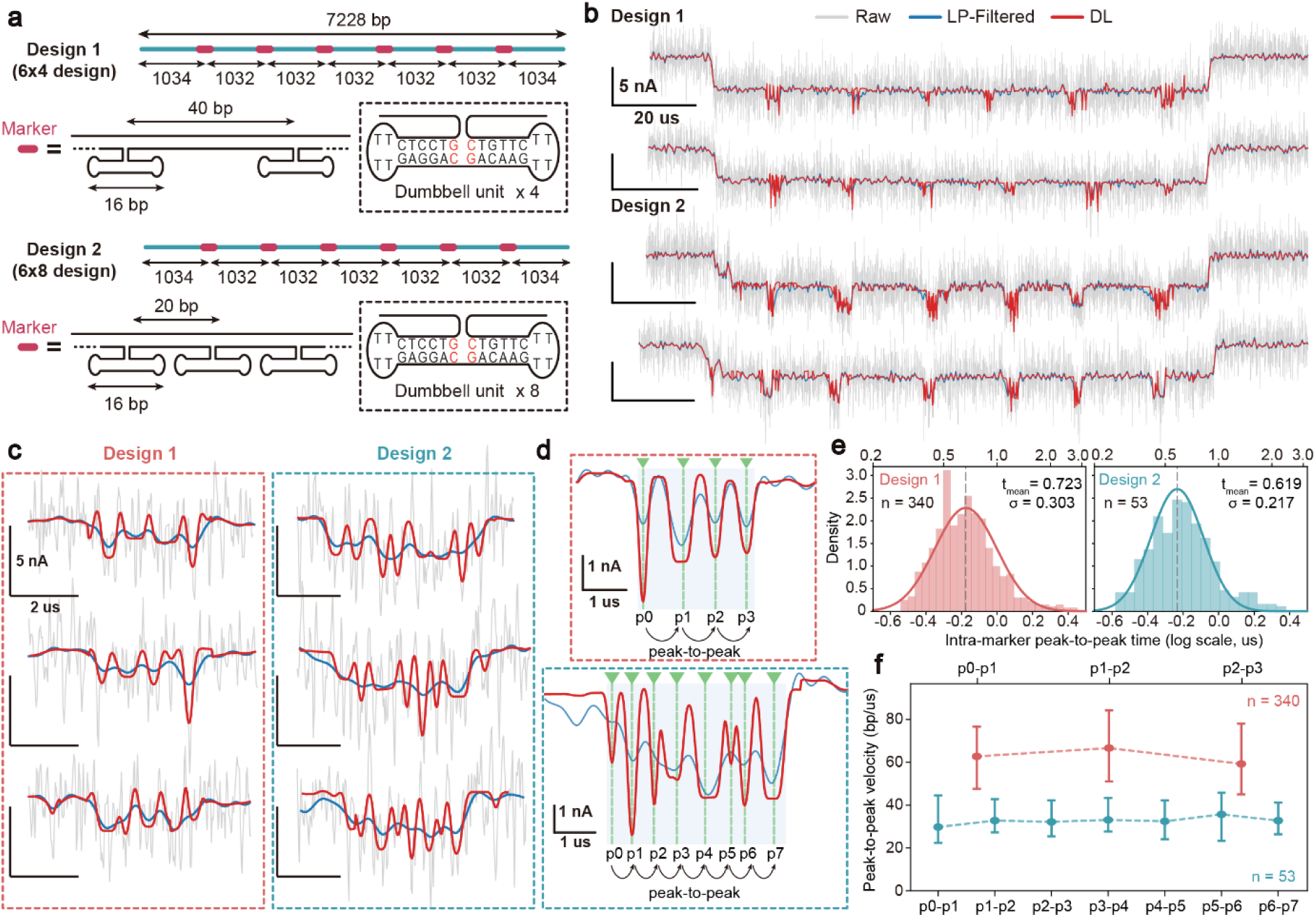
Detection of a few nanometer-spaced DNA nanostructures with sub-microsecond temporal resolution. **(a)** Schematic plots of two DNA molecule designs carrying different amounts of dumbbell units. Both designs have 6 identical marker regions while different dumbbell density in the markers, with design 1 containing 4 dumbbells per marker and design 2 containing 8 per marker. Two C-G pairs are added in the original sequence of individual dumbbell units. **(b)** Representative translocation events of the two DNA molecules in 3 M LiCl solution through a 7 nm-in diameter pore under 1000 mV voltage bias. All three traces are superimposed on same events, including the raw trace (in grey) collected under 10 MHz bandwidth and 40 MHz sampling rate, low-pass (LP)-filtered trace (in blue) with cutoff frequency at 2 MHz, and deep-learning (DL) reconstructed signal (in red). **(c)** Zoomed-in view of translocation signals in marker and dumbbell region, with same colour indication as in **(b)**. **(d)** Examples of peak detection results (green dashed line) from the DL reconstructed signal of two different molecules. Each detected peak is defined with an id and peak-to-peak time interval is defined accordingly using all peak pairs. **(e)** The intra-marker peak-to-peak time distribution of two DNA molecule designs (n = 340 for Design 1 and n = 53 for Design 2) are plotted in log-scale and fitted with Gaussian (solid lines). Bottom and top axes are in logarithmic and linear scale, respectively. **(f)** Intra-marker peak-to-peak velocity profile. The velocities are calculated using centre-to-centre distances of 40 bp for Design 1 and 20 bp for Design 2 and corresponding peak-to-peak time interval for each neighbouring pair of the peaks inside the marker, and statistically summarized across events. The error bar represents the standard deviation. Colour representation in **(c-f)**: Design 1 in magenta, Design 2 in cyan.

The representative translocation event traces from these two designs showed similar structure of current blockage profile as previous sample, including a primary level representing dsDNA backbone and six secondary levels caused by the extruded markers on the DNA backbone (**Figure 4b**). For both designs, the DL network recovers the expected multi-peak patterns within individual markers, whereas the corresponding LP-filtered trace likely merges closely spaced features or fragment those dumbbell peaks inconsistently depending on the applied digital filter (**Figure 4c**). The clearly resolved intra-marker signals from DL reconstruction allowed us to statistically analyse the peak spacing. As illustrated in **Figure 4d**, detected peaks are labelled with an id from p0 to p3 in the 4-dumbbell-per marker design (design 1) and from p0 to p7 for the 8-dumbbell-per-marker design (design 2). As the designed dumbbell centre-to-centre distance inside a marker is constant, all neighbouring pairs of detected peaks (p0-p1, p1-p2, p2-p3, etc.) ideally represent the same physical distance of dumbbells. We therefore plotted the distribution of intra-marker peak-to-peak time intervals for the two DNA constructs and extracted the mean intervals with Gaussian fitting (**Figure 4e**). The slightly higher mean peak-to-peak interval of 0.723 µs in design 1 corresponds to the larger spacing between dumbbells. Likely, the mean interval of 0.619 µs in the design 2 is similar to the statistical result in the 2-marker design, which has nearly the same dumbbell sidearm spacing as in design 2. The peak-to-peak intervals remain broadly consistent across successive dumbbell pairs within each design, indicating that the reconstructed signal retained the physical barcode geometry rather than generating arbitrary local features.

Moreover, benefitting from the clearly reconstructed peak inside the marker, we computed for the first time from experimental data the dumbbell-to-dumbbell velocity profiles inside individual markers and explored its dependence on dumbbell density (**Figure 4f**). The results show that average intra-marker molecule free translocation velocity is 62.8-68.7 bp/µs for design 1 and 34.7-38.5 bp/µs for design 2, while the denser marker with more dumbbells present slower translocation velocity. Similar trends have been observed while calculating the fractional contour velocity profile of the entire translocation event, as the molecular average velocity in the marker region is slightly smaller than the inter-marker velocity (**Supplementary Fig.16**). This is likely due to the increased pore-polymer interactions in the marker region where the molecule presents larger cross-sectional areas, as reported in a previous study^45^. These results on various designs further demonstrated the capability of our hardware-software co-designed platform in recovering nm-spaced barcode units from high-speed solid-state nanopore recording and can therefore extend the usable information content of fast translocation events.

### Transferability test via dual-channel recording

To assess whether the platform is transferable across distinct readout chains and support potential application in multi-channel high-throughput nanopores sensing system, we performed simultaneous recordings of the same nanopore translocation events using two independent channels connected to a single low-noise nanopore device (**Figure 5a**). This configuration provided a stringent validation test as the two channels share the same physical event while exhibiting intrinsically different noise baselines. A 6.3 nm-in diameter nanopore is measured in 4 M LiCl (pH 9.0) solution using a 10 MHz-bandwidth multi-channel amplifier to simulate a different measurement condition from previous single-channel measurement. All three DNA constructs were detected with a cross-membrane driving voltage of 1000 mV via a differential of 500 mV on CH1 and −500 mV on CH2. Recorded translocation events were processed with a single DL model re-trained on the same architecture while adapted for a different real-life noise characteristic of the amplifier.

**Figure 5.**
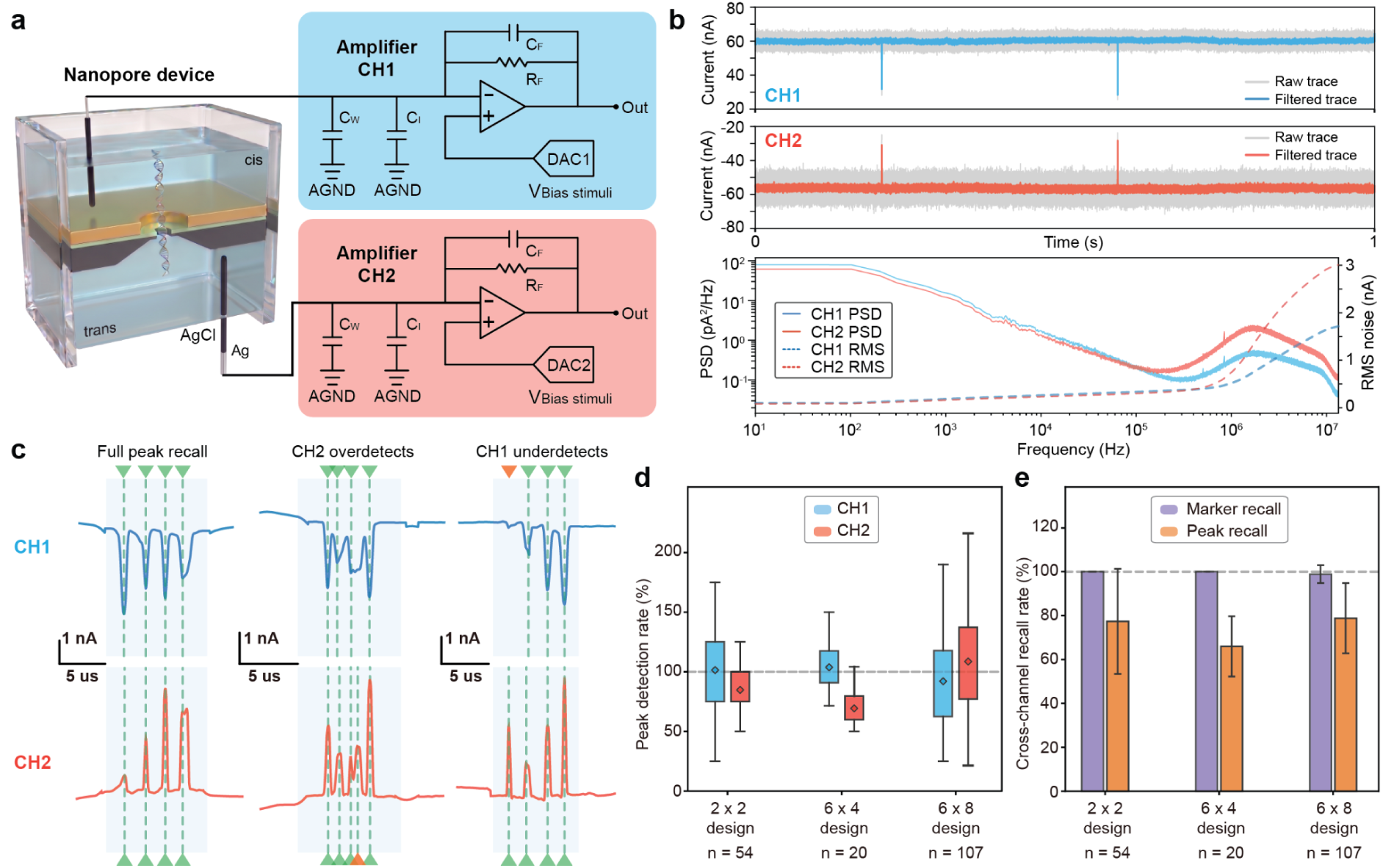
Transferability across dual-channel recording on a single nanopore device. **(a)** Schematic of the dual-channel measurement configuration. Two-independent voltage-clamp amplifiers (CH1 and CH2) are connected respectively to the *cis* and *trans* chambers of the same low-noise nanopore device, with independent DAC control of the applied biases. Same DNA translocation signals from the nanopore are recorded simultaneously by two readout channels. **(b)** Representative synchronized ionic current traces of translocation events on the two channels (top) and baseline current PSD and RMS noise of each channel (bottom). The current trace is from a single 6.3-nm nanopore in 4 M LiCl (pH 9.0) using a 10 MHz-bandwidth multi-channel amplifier at 26 MHz sampling frequency, with applied bias of 500 mV on CH1 and −500 mV on CH2. Raw traces (in grey) are overlaid with LP-filtered trace (in blue for CH1, in red for CH2) with cutoff frequency of 2 MHz. **(c)** Examples of cross-channel peak recalling in matched marker regions from 6×4 DNA construct, based on DL reconstructed results. Detected peaks are marked with green dashed lines. Cross-channel peak recalling is marked with triangle shapes, with recalled in green and mis-matched in orange. **(d)** Mean peak-detection rate based on DL reconstructed signal in each channel for each DNA construct, calculated by detected peak in each event over designed peak amounts. Detailed values in Supplementary Table 6. **(e)** Cross-channel marker and dumbbell-peak recall rates for all three DNA constructs. Both recall rates are defined as whether corresponding marker or peak is detected in CH2 when the marker or peak is detected in CH1. Detailed values in Supplementary Table 7.

Due to the differential voltage configuration, the typical ionic current traces from dual-channel measurement have opposite polarity, as well as the direction of current blockage spikes (**Figure 5b**). The two amplifier channels present different high-frequency noise characteristics as a result of the configuration because the voltage bias is applied from the active input side instead of reference side, which is usually adopted for single-channel measurement. Such conditions therefore provided the scenario of having independent readout channels with distinct noise characteristics, which is ideally expected to avoid while inevitable in multi-channel high-throughput measurements. Moreover, we also simulated different measurement conditions by varying the buffer solution. The model is re-trained based on the measured noise characteristics in the completely different environment. We reconstructed the two channel signals using a neural network with the same set of weights, and the noise level used for the synthesized data in the training process was the average of the two channels.

The cross-channel feature recall rate helps evaluate the transferability of the platform, which indicates the percentage of detected features such as marker or peak in one channel that is recovered in the other channel. **Figure 5c** shows representative examples of the peak recalling results across the two channels from 6×4 DNA constructs. Each marker is first detected from the respective channel and then paired accordingly to the temporal window after channel synchronization, followed by peak recalling inside individual marker regions across the two channels based on DL reconstructed signals. We found that either full peak recall or peak missing and overdetection could happen (**Figure 5c,d**), likely due to the very distinct noise baselines of the two channels here and the difference of DL model sensitivity on each channel. On the other hand, it also shows that the dual-channel measurement system could potentially increase the average detection accuracy by comparing and cross-validating results from individual channels, and such accuracy increment is expected for future multi-channel measurement. A statistical analysis of cross-channel recall rates shows almost 100% recalling of marker regions and on average 65-80% recalling of peaks inside paired markers (**Figure 5e**), which we expect to be higher with reduced cross-channel noise baseline variations. The result therefore demonstrates the good consistency of DL reconstruction on the high-speed translocation signals and general transferability of the measurement platform.

## Discussion

In this study, we developed a transferable high-bandwidth solid-state nanopore sensing platform that combines wafer-scale dielectric engineering with deep-learning-based signal reconstruction method to recover closely spaced barcode features on fast translocating molecules. With the integration of a low-dielectric SU8 coating layer on a suspended silicon nitride nanopore chip, we substantially reduced device capacitance to pF level and suppressed high-frequency electrical noise in a scalable and reproducible way, resulting in 5-fold reduction of nanopore noise compared with bare SiN_x_ membrane. This improvement brings SiN_x_ membrane-based nanopore noise performance close to glass-substrate nanopores while keeping the fabrication flexibility and integration potential of silicon-based devices. This hardware advance expanded the usable temporal bandwidth of the measurement system and enabled direct recording of fast translocation events at 40 MHz sampling rate.

Leveraging on the improved acquisition bandwidth, we designed a deep-learning reconstruction network to recover translocation blockage profile and closely spaced sublevels present in the noisy raw trace. Using engineered DNA dumbbell barcodes as a benchmark, the platform resolved nanometer-scale structural features with sub-microsecond temporal resolution, and allowed us to experimentally quantify the intra-marker spacing and velocity trends across various constructs. The combined platform therefore demonstrates the capability to uncover meaningful structural information from fast translocating molecules. With dual-channel recording of translocation events on a single nanopore device, we further showed that the hardware and software co-designed platform is transferable across independent acquisition channels with distinct baseline noise characteristics. This transferability is essential for future multichannel nanopore measurements where cross-device, cross-channel and operating conditions could vary.

The present study highlights the importance of integrating device engineering and computational reconstruction into a unified strategy for high-speed nanopore measurements. It establishes a general route to recover structurally informative features from rapid translocation events that would otherwise remain inaccessible with insufficient acquisition bandwidth and conventional signal processing approaches. This platform could be potentially extended to denser and smaller molecular barcodes to support DNA structure-based data storage or to other biomolecular targets such as proteins. With further integration into multi-channel readout schemes based on the wafer-scale fabrication compatibility, the same working principle may help realize reproducible high-throughput and more information-rich single-molecule sensing across diverse device, readout and experimental conditions.

## Methods

### Nanopore fabrication

The silicon nitride (SiN_x_) nanopore devices used in this study were fabricated in wafer-scale, starting from the standard 4-inch Boron-doped thick silicon wafers (Si, 380 ± 10 µm in thickness, with 60 nm dry silicon oxide (SiO_2_) layer and a 20 nm low-stress silicon nitride (SiN_x_) layer on both sides, supplied by EPFL Center of MicroNano Technology (CMi)). Multiple photolithography and dry etching processes were done in the CMi cleanroom facility to form a suspending SiN_x_ membrane (40 µm x 40 µm in size, 20 nm-thick) on the front side of the wafer, as described in previous work^46^. The wafer was cleaned in acid piranha solution overnight to remove contamination from photolithography. After the deionised water rinsing, drying, and a short baking process at 300 ℃, the wafer was ready for the SU8 coating to reduce the noise. To improve the coating uniformity of the SU8 layer on the SiN_x_ surface, the wafer was treated in oxygen plasma. An SU8 (Kayaku Advanced Materials Inc.) layer was immediately spin-coated on the hydrophilic wafer surface. Thickness of the SU8 layer was tuned via spin-coating velocity, and window size of SU8 apertures were modulated by varying the design pattern used in the direct-laser writing photolithography process (**Supplementary Information Section 1**). Each chip was then cleaved manually out from the wafer, and the nanopore was produced in a well-controlled way through transmission electron microscope (TEM) drilling on the SiN_x_ membrane.

### Nanopore electrical noise characterisation

The SU8-coated SiN_x_ nanopore chip was treated with oxygen plasma to promote surface wettability prior to assembly in a customised flowcell (Elements srl, Italy). Both *cis* and *trans* chambers of the flowcell were filled with buffered solution of 1 M KCl, 10 mM Tris-HCl (pH 8.0) and 1 mM EDTA. By applying a 100 mV voltage bias on the *trans* chamber, ionic currents were recorded using Nanopore Reader 10 MHz (Elements srl, Italy) with 10 MHz bandwidth and 40 MHz sampling rate. The power spectral density (PSD) and root-mean-square (RMS) current noise were computed from the acquired high-bandwidth traces. The total system capacitance of each nanopore device was determined using a triangle-wave voltage stimulation protocol. A symmetrical triangular waveform was applied with frequency of 5 Hz and wave amplitude at 100 mV (corresponding to voltage sweep rate *α* = 2000 *V*/*s*), and resulting current responses were analysed to extract the capacitance information. The 4 mm x 4 mm glass-substrate nanopore chips were purchased commercially (Norcada, Edmonton, AB, Canada) with SiN_x_ membrane of around 35 µm in diameter, and nanopores were drilled in TEM. Further details of noise characterisation are provided in **Supplementary Information Section 1**. The ionic current noise measured under different electrolyte solutions (1M KCl, 3M LiCl and 4M LiCl) are characterised, as shown in **Supplementary Fig.17**.

### Synthesis of DNA with nanostructures

The DNA molecules with nanostructure markers were prepared by hybridising a linearised M13mp18 single-stranded DNA scaffold together with different sets of complementary oligonucleotide mixtures. Circular single-stranded M13mp18 plasmid (New England Biolabs, MA, USA) was first linearised by restriction enzyme cutting at BamHI and EcoRI sites. In total 190 short oligonucleotides were designed to complement the single-stranded scaffold as in previous work^28^. The different types of DNA molecules with dumbbell nanostructures were formed by attaching certain oligonucleotides with additional DNA structure at designed positions. The sequence details of all the strands and synthesis protocols are provided in **Supplementary Information Section 3**.

### Single-channel nanopore measurement

For single-channel recordings, all measurements were performed in 3 M LiCl buffer solution (pH 8.0), with 10 mM Tris-HCl and 1 mM EDTA. The low-noise nanopore devices were prepared and baseline current was checked for blank control. The DNA samples were diluted to a final concentration of ∼1 nM using the measurement buffer solution and then added to the *cis* side of the nanopore chip. Positive voltage biases from 500 mV and 1000 mV were applied on the *trans* side to drive molecules to translocate through the pore. The ionic current signals were recorded using a Nanopore Reader 10 MHz system (Elements srl, Italy) at 10 MHz bandwidth and 40 MHz sampling rate. High-bandwidth time-series ionic current signals were first processed with commercial software Nanolyzer (Northern Nanopore Instruments, Canada), and DNA translocation events were screened out from digitally low-pass filtered data with filter cutoff frequency at 2 MHz. After the event detection, 40 MHz high-bandwidth raw data was used for subsequent analysis.

### Multi-channel nanopore measurement

Single low-noise nanopore device was assembled in the customised flowcell and the *cis* and *trans* chambers were connected respectively to two active channels CH1 and CH2 on the front-end of Nanopore Reader 4×10 MHz instrument (Elements srl, Italy), enabling simultaneous voltage bias stimulus and synchronised ionic current signal recordings via two individual amplifiers. A different buffer solution of 4 M LiCl, 10 mM Tris-HCl and 1 mM EDTA at pH 9.0 was used for the multi-channel measurements to simulate different experimental conditions. The measurement buffer was filled in both chambers of the flowcell and ionic current signals were recorded on both channels after applying a differential voltage bias across the nanopore membrane (250 mV for CH1 and −250 mV for CH2 for a differential voltage of 500 mV, for example). DNA samples were similarly diluted with the buffer solution to ∼1 nM and then added to the *trans* chamber. The ionic current signals for molecule translocation were acquired at 10 MHz bandwidth and 26 MHz sampling rate. Event-specific segments of the signal were extracted first with a 2 MHz low-pass digital filter on both channels. Different event sets were detected through each channel due to different current baselines. Custom analysis was therefore applied to correlate translocation events across two channels and high-bandwidth raw data of the correlated events were used for subsequent analysis.

### Network Training

#### Noise Characterisation and Synthetic Data Generation

Prior to training, we characterised the noise spectral properties of the experimental ionic current recordings using PSD analysis. The noise was decomposed into four physically motivated components based on its frequency signature: (i) pink (1/f) noise, arising from low-frequency baseline drift; (ii) white noise, with a flat power spectrum; (iii) dielectric noise, with power scaling as *f*¹, attributed to dielectric loss from the chip; and (iv) capacitive noise, with power scaling as *f*², caused by coupling of voltage noise through the input capacitance of the recording amplifier and the device. For each component, the RMS noise was estimated by integrating the PSD within the corresponding frequency band.

A full synthetic training dataset was built on measured real-life noise. Each training trace was generated in four steps: (1) Event generation. A clean, less-noisy ionic current signal was constructed as a piecewise-constant trace (with multiple rectangular blockade events) with length of 2048 data points. The number and depth of event blockade on each trace was randomly generated. (2) Edge softening. The clean signal was convolved with a zero-phase 8th-order Bessel low-pass filter at a randomly sampled cutoff ratio of 1–10% of the sampling frequency, simulating the bandwidth limitation of the recording amplifier. This step can smooth out the sharp edges of the ideal event, reducing the ringing effect in the network fitting and constitutes the training target (ground truth label) in network training. (3) Noise mixing. Each of the four noise components was synthesized at normalized variance and scaled by independently drawn random amplitudes. The four weighted noise signals were then summed and added to the bandwidth-limited clean signal to produce the noisy input. (4) Multi-scale filter input. The noisy signal was processed through a bank of zero-phase 8th-order Bessel low-pass filters at multiple cutoff ratios, plus the unfiltered raw signal, yielding a multi-scale input stack. All channels were z-score normalized using the median and standard deviation of the unfiltered raw channel, preserving inter-channel relative amplitude differences that encode frequency content.

#### Network Architecture

We designed a 1D U-Net adapted from the original 2D biomedical image segmentation architecture for one-dimensional ionic current signals^47^. The encoder–decoder structure naturally captures multi-scale temporal features through successive downsampling and upsampling, with skip connections preserving fine-grained edge detail. There are three structures in the architecture. (1) Encoder. The network begins with an initial feature extraction block that maps the multi-scale data input to 32 feature channels. Three successive encoder stages apply max-pooling followed by a residual convolutional block, progressively doubling the feature dimension: 32 → 64 → 128 → 256 (bottleneck). (2) Residual convolutional blocks. Each block consists of three sequential 1D convolutions, each followed by Batch Normalization. The SiLU function is applied as the activation. A residual shortcut is added before the final SiLU, forming a pre-activation residual connection that stabilizes gradient flow in deep networks. (3) Decoder and Output. Three convolution layers progressively upsample the bottleneck representation back to the original signal length. At each decoder level, the upsampled feature map is concatenated with the corresponding encoder skip connection, and processed by another residual block, following the standard U-Net skip fusion pattern. The decoder reduces channels in the reverse order: 256 → 128 → 64 → 32. A final 1×1 convolution maps the feature map to a single-channel denoised signal prediction.

#### Loss Functions

Training used an edge-preserving composite loss:

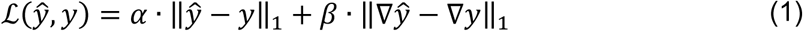

Where *ŷ* is the network prediction, *y* is the ground truth, ∇ denotes the first discrete difference (temporal gradient). The first term represents fidelity, which measures the difference in amplitude between the predicted and the actual values. The second is the edge preserving item that averages the absolute gradient error, where the gradient refers to the first-order difference between adjacent data points. This term encourages predictions to have similar local variation trends to the non-noisy signal, thereby preserving edge structures.

#### Training Protocol

The model was trained using the AdamW optimizer (learning rate 1 × 10^−4^, default weight decay) for 300 epochs with a batch size of 128. Training data were generated on-the-fly by 8 parallel workers (each with an independent random seed derived from the PyTorch initial seed), ensuring that each epoch saw a different random realization of event positions, noise amplitudes, and noise colours. All experiments were conducted on a computer with RTX3090 and 64 GB RAM.

### Data Analysis

#### Single-channel data processing

Raw ionic current traces from all three DNA constructs (2×2, 6×4 and 6×8 designs) were acquired at 40 MHz sampling rate and pre-processed using Poriscope^48^ to extract translocation events associated with dumbbell-containing DNA molecules. Event identification was performed on 2 MHz Bessel low-pass filtered trace using a thresholding-crossing algorithm with an SNR of 10. Both raw and low-pass filtered data of each detected event were then stored individually for downstream analysis. The translocation event data were processed through a two-stage detection pipeline (**Supplementary Information Section 4**). In the first stage, the deep learning (DL) denoiser reconstructed a clean signal from each translocation event, suppressing high-frequency noise while preserving sub-microsecond dumbbell features. Coarse marker-region boundaries were identified on the 2 MHz Bessel-filtered reference signal using prominence- and width-constrained peak detection, localising each dumbbell-cluster region along the translocation trace. In the second stage, individual dumbbell peak positions were resolved within each search window using the DL-reconstructed signal, preventing noise artifacts outside the marker-boundaries from contributing false positive detections. Peak prominences were evaluated relative to the pre-event baseline noise estimated by the standard deviation of a Gaussian distribution. Events were retained for statistical analysis when the number of detected marker regions matched the expected design geometry. More examples of peak detection results from all three DNA constructs are shown in **Supplementary Fig.18-20**. Detected peak-to-peak times were converted to translocation velocities using the known centre-to-centre distances of adjacent dumbbell pairs, and inter-peak spacing distributions were characterised by log-normal fits in linear space and Gaussian fits in log₁₀ space to extract median velocities and geometric standard deviations.

#### Dual-channel data processing

Raw ionic current traces recorded at 26 MHz sampling frequency were pre-processed to extract and then pair the events from each channel. Due to the different baseline, the threshold-based event detection resulted in different event sets detected from each channel. Corresponding events were paired using a greedy 1-to-1 matching algorithm based on temporal overlap, meaning that candidate pairs were scored by the intersection over union (IoU) of their time spans, and assignments were made in descending IoU score to retain the paired event. The paired events were then passed to the DL denoiser. The model was re-trained on the same architecture with different real-life noise collected from the amplifier. Prior to marker and peak detection, each paired event was temporally aligned to account for the phase shift between two independent amplifier channels. The integer-sample lag between channels was determined by cross-correlation of the 2 MHz Bessel-filtered signals, and all the raw, filtered and DL-reconstructed traces were sliced to their shared overlapping window to synchronize. Subsequently, a similar two-stage detection pipeline is used as single-channel data analysis to locate the marker windows and then detect dumbbell peaks based on DL-reconstructed signal in each channel. Peak detection rate was defined as the number of detected peaks normalized by the expected count per event according to the construct design. Cross-channel peak matching was performed on DL-reconstructed signal by pairing each peak detected on channel 1 with the temporally nearest peak on channel 2. Cross-channel recall was defined as the fraction of channel-1 peaks that received a match (n_matched / n_peaks_ch1 × 100%). Marker consistency was quantified per event as min(n_markers_ch1, n_markers_ch2) / n_markers_ch1 × 100%, reflecting the degree to which both channels resolved the same number of marker windows and supporting that the peak recall rate was estimated within matched markers instead of false positive peaks outside the marker region.

## Data availability

Source data for each figure in this paper are provided, including raw data of ionic current traces for noise characterisation, translocation events and deep-learning reconstructed current signals.

## Code availability

The code used for neural network construction and training will be available before publication.

## Acknowledgement

The authors acknowledge the financial support from the Swiss National Science Foundation through the National Centre of Competence in Research Bio-Inspired Materials and Grant No. 200021_192037. The authors thank the EPFL Center of MicroNanoTechnology (CMi) for help with chip fabrication and characterisation, and the EPFL Center for Electron Microscopy (CIME) for access to electron microscopes.

## Author Information

### Authors and Affiliations

**Laboratory of Nanoscale Biology, Institute of Bioengineering, Ecole Polytechnique Fédérale de Lausanne (EPFL), Lausanne, Switzerland**

Nianduo Cai, Wei Guo, Yunfei Teng, Akhil Sai Naidu, Tzu-Heng Chen, Aleksandra Radenovic

**NCCR Bio-inspired Materials, École Polytechnique Fédérale de Lausanne, Lausanne, Switzerland**

Yunfei Teng, Aleksandra Radenovic

**Programmable Biomaterials Laboratory, Institute of Materials, Ecole Polytechnique Fédérale de Lausanne (EPFL), Lausanne, Switzerland**

Yameng Lou, Siu-Ho Wong, Maartje Bastings

**Elements Srl, Cesena, Italy**

Filippo Cona, Federico Thei

**Department of Materials Science and Engineering, National Yang Ming Chiao Tung University (NYCU), Hsinchu, Taiwan (R.O.C)**

Tzu-Heng Chen

## Contributions

N.C., T.-H. C., and A.R. conceived the idea and designed the study. Y.T. fabricated the SiN_x_ nanopore substrates. N.C. designed and fabricated the low-noise nanopore devices. N.C. and Y.T. performed TEM characterisations of the nanopore device. Y.L., S.-H. W. and N.C. designed and validated the DNA constructs. Y.L. synthesized the DNA constructs and characterised with gel electrophoresis. M.B. instructed on the design and the preparation of DNA constructs. N.C. and T.-H. C. performed the electrical noise characterisation experiments. N.C. performed all DNA translocation experiments. F.C. and F.T. configured the dual-channel instrument and instructed on measurement. W.G. developed the idea of deep-learning denoiser, trained the neural networks and processed the data with the deep-learning model. N.C. performed data analysis. N.C. and A.R. wrote the first draft of the manuscript. A.R. supervised the work. All authors contributed to the writing and discussion of the manuscript.

## References

1. Venkatesan, B. M. & Bashir, R. Nanopore sensors for nucleic acid analysis. Nature Nanotech 6, 615–624 (2011).

2. Laszlo, A. H. et al. Decoding long nanopore sequencing reads of natural DNA. Nat Biotechnol 32, 829–833 (2014).

3. Yusko, E. C. et al. Controlling protein translocation through nanopores with bio-inspired fluid walls. Nature Nanotech 6, 253–260 (2011).

4. Storm, A. J. et al. Fast DNA Translocation through a Solid-State Nanopore. Nano Lett. 5, 1193–1197 (2005).

5. Kowalczyk, S. W., Hall, A. R. & Dekker, C. Detection of Local Protein Structures along DNA Using Solid-State Nanopores. Nano Lett. 10, 324–328 (2010).

6. Wanunu, M. et al. Rapid electronic detection of probe-specific microRNAs using thin nanopore sensors. Nature Nanotech 5, 807–814 (2010).

7. Bell, N. A. W. & Keyser, U. F. Digitally encoded DNA nanostructures for multiplexed, single-molecule protein sensing with nanopores. Nature Nanotech 11, 645–651 (2016).

8. Smeets, R. M. M., Kowalczyk, S. W., Hall, A. R., Dekker, N. H. & Dekker, C. Translocation of RecA-Coated Double-Stranded DNA through Solid-State Nanopores. Nano Lett. 9, 3089–3095 (2009).

9. Plesa, C. et al. Fast Translocation of Proteins through Solid State Nanopores. Nano Lett. 13, 658–663 (2013).

10. Fologea, D., Uplinger, J., Thomas, B., McNabb, D. S. & Li, J. Slowing DNA Translocation in a Solid-State Nanopore. Nano Lett. 5, 1734–1737 (2005).

11. Wanunu, M., Sutin, J., McNally, B., Chow, A. & Meller, A. DNA Translocation Governed by Interactions with Solid-State Nanopores. Biophysical Journal 95, 4716–4725 (2008).

12. Manrao, E. A. et al. Reading DNA at single-nucleotide resolution with a mutant MspA nanopore and phi29 DNA polymerase. Nat Biotechnol 30, 349–353 (2012).

13. Olasagasti, F. et al. Replication of individual DNA molecules under electronic control using a protein nanopore. Nature Nanotech 5, 798–806 (2010).

14. Cherf, G. M. et al. Automated forward and reverse ratcheting of DNA in a nanopore at 5-Å precision. Nat Biotechnol 30, 344–348 (2012).

15. Derrington, I. M. et al. Nanopore DNA sequencing with MspA. Proceedings of the National Academy of Sciences 107, 16060–16065 (2010).

16. Wang, C., Sensale, S., Pan, Z., Senapati, S. & Chang, H.-C. Slowing down DNA translocation through solid-state nanopores by edge-field leakage | Nature Communications. Nature Communications 12, 140 (2021).

17. Larkin, J. et al. Slow DNA Transport through Nanopores in Hafnium Oxide Membranes. ACS Nano 7, 10121–10128 (2013).

18. Soni, N., Chandra Verma, N., Talor, N. & Meller, A. Over 30-Fold Enhancement in DNA Translocation Dynamics through Nanoscale Pores Coated with an Anionic Surfactant. Nano Lett. 23, 4609–4616 (2023).

19. Chou, Y.-C. et al. Coupled nanopores for single-molecule detection. Nat. Nanotechnol. 19, 1686–1692 (2024).

20. Kowalczyk, S. W., Wells, D. B., Aksimentiev, A. & Dekker, C. Slowing down DNA Translocation through a Nanopore in Lithium Chloride. Nano Lett. 12, 1038–1044 (2012).

21. Smeets, R. M. M. et al. Salt Dependence of Ion Transport and DNA Translocation through Solid-State Nanopores. Nano Lett. 6, 89–95 (2006).

22. Dekker, C. Solid-state nanopores. Nature Nanotech 2, 209–215 (2007).

23. Zeng, S., Wen, C., Solomon, P., Zhang, S.-L. & Zhang, Z. Rectification of protein translocation in truncated pyramidal nanopores. Nat. Nanotechnol. 14, 1056–1062 (2019).

24. Kwok, H., Briggs, K. & Tabard-Cossa, V. Nanopore Fabrication by Controlled Dielectric Breakdown. PLOS ONE 9, e92880 (2014).

25. Rosenstein, J. K., Wanunu, M., Merchant, C. A., Drndic, M. & Shepard, K. L. Integrated nanopore sensing platform with sub-microsecond temporal resolution. Nat Methods 9, 487–492 (2012).

26. Fragasso, A., Schmid, S. & Dekker, C. Comparing Current Noise in Biological and Solid-State Nanopores. ACS Nano 14, 1338–1349 (2020).

27. Balan, A. et al. Improving Signal-to-Noise Performance for DNA Translocation in Solid-State Nanopores at MHz Bandwidths. Nano Lett. 14, 7215–7220 (2014).

28. Chen, K. et al. Super-Resolution Detection of DNA Nanostructures Using a Nanopore. Advanced Materials 35, 2207434 (2023).

29. Chen, K. et al. Dynamics of driven polymer transport through a nanopore | Nature Physics. Nature Physics 17, 1043–1049 (2021).

30. de Vreede, L. J. et al. Wafer-scale fabrication of fused silica chips for low-noise recording of resistive pulses through nanopores. Nanotechnology 30, 265301 (2019).

31. Dimitrov, V. et al. Nanopores in solid-state membranes engineered for single molecule detection. Nanotechnology 21, 065502 (2010).

32. Ma, Q. D. Y. et al. Robust Noise Separation and Denoising of Ionic Current Signals for Nanopore Sensors Based on Independent Component Analysis. J. Phys. Chem. Lett. 16, 10612–10620 (2025).

33. Tsutsui, M., Takaai, T., Yokota, K., Kawai, T. & Washio, T. Deep Learning-Enhanced Nanopore Sensing of Single-Nanoparticle Translocation Dynamics. Small Methods 5, 2100191 (2021).

34. Taniguchi, M. et al. Combining machine learning and nanopore construction creates an artificial intelligence nanopore for coronavirus detection. Nat Commun 12, 3726 (2021).

35. Zhang, M. et al. Real-time detection of 20 amino acids and discrimination of pathologically relevant peptides with functionalized nanopore. Nat Methods 21, 609–618 (2024).

36. Wang, K. et al. Unambiguous discrimination of all 20 proteinogenic amino acids and their modifications by nanopore. Nat Methods 21, 92–101 (2024).

37. Dematties, D., Wen, C., Pérez, M. D., Zhou, D. & Zhang, S.-L. Deep Learning of Nanopore Sensing Signals Using a Bi-Path Network. ACS Nano 15, 14419–14429 (2021).

38. Misiunas, K., Ermann, N. & Keyser, U. F. QuipuNet: Convolutional Neural Network for Single-Molecule Nanopore Sensing. Nano Lett. 18, 4040–4045 (2018).

39. Smeets, R. M. M., Keyser, U. F., Dekker, N. H. & Dekker, C. Noise in solid-state nanopores. Proceedings of the National Academy of Sciences 105, 417–421 (2008).

40. Tabard-Cossa, V., Trivedi, D., Wiggin, M., Jetha, N. N. & Marziali, A. Noise analysis and reduction in solid-state nanopores. Nanotechnology 18, 305505 (2007).

41. Wen, C. et al. Generalized Noise Study of Solid-State Nanopores at Low Frequencies. ACS Sens. 2, 300–307 (2017).

42. Zhang, Y. et al. Rapid detection of neurons in widefield calcium imaging datasets after training with synthetic data. Nat Methods 20, 747–754 (2023).

43. Lu, Z. et al. Virtual-scanning light-field microscopy for robust snapshot high-resolution volumetric imaging. Nat Methods 20, 735–746 (2023).

44. Yuan, Z., Liu, Y., Dai, M., Yi, X. & Wang, C. Controlling DNA Translocation Through Solid-state Nanopores. Nanoscale Research Letters 15, 80 (2020).

45. Charron, M., Elliott, B., Kerrouri, N., He, L. & Tabard-Cossa, V. Velocity Fluctuation and Force Scaling During Driven Polymer Transport through a Nanopore. ACS Nano 19, 30950–30960 (2025).

46. Cai, N., Chen, T., Teng, Y., Naidu, A. S. & Radenovic, A. Nanopore Trap for Label-Free Fingerprinting of Surface-modified Single Nanoparticles. Small Methods 10, e01765 (2026).

47. Ronneberger, O., Fischer, P. & Brox, T. U-Net: Convolutional Networks for Biomedical Image Segmentation. Preprint at 10.48550/arXiv.1505.04597 (2015).

48. González González, A. C., Kerrouri, N., Wadhwa, D., Tabard-Cossa, V. & Briggs, K. Poriscope: A Configurable Pipeline for Nanopore Data Analysis. Journal of Open Research Software 14, (2026).

