## Supplementary Information for "A generalizable codesigned platform for solid-state nanopore sensing beyond the capacitive-noise constraints"

### Table of Contents

**Section 1. Wafer-scale fabrication of low-noise nanopore devices and noise characterizations**

**Section 2. Design of deep learning framework to recover embedded transient feature in high-bandwidth nanopore translocation signal**

**Section 3. Design and synthesis of DNA molecules carrying dumbbell nanostructures**

**Section 4. Nanopore measurement and data analysis**

**References**

#### Section 1. Wafer-scale fabrication of low-noise nanopore devices and noise characterizations

##### 1.1 Wafer-scale fabrication process flow

The low-noise chips are fabricated on 4-inch silicon wafers through a wafer-scale microfabrication process (Figure S1.a). A 60 nm $SiO_2$ layer is first thermally grown on both sides of the silicon substrate, followed by deposition of 20 nm low-stress $SiN_x$ layers by LPCVD. Backside patterns defining membrane windows and cleaving marker lines are generated by direct laser writing, followed by reactive ion etching of the $SiN_x/SiO_2$ layers to transfer the pattern on the wafer. Suspended $SiN_x$ membranes are subsequently formed through anisotropic wet etching of the silicon substrate using KOH solution. A 50 μm SU8 layer is then spin-coated on the front side $SiN_x$ membrane and patterned by mask-less photolithography to define apertures above the suspending membrane region. The yield per wafer depends on the backside pattern, and with the design shown in Figure S1.b, 204 devices are fabricated in parallel on a single 4-inch wafer, which could potentially be further increased with a tighter layout and smaller single-chip size. The SU8 coating on the frontside membrane also improves the robustness of the device via covering most of the membrane surface (Figure S1.c) and protecting the thin $SiN_x$ window from mechanical damage.

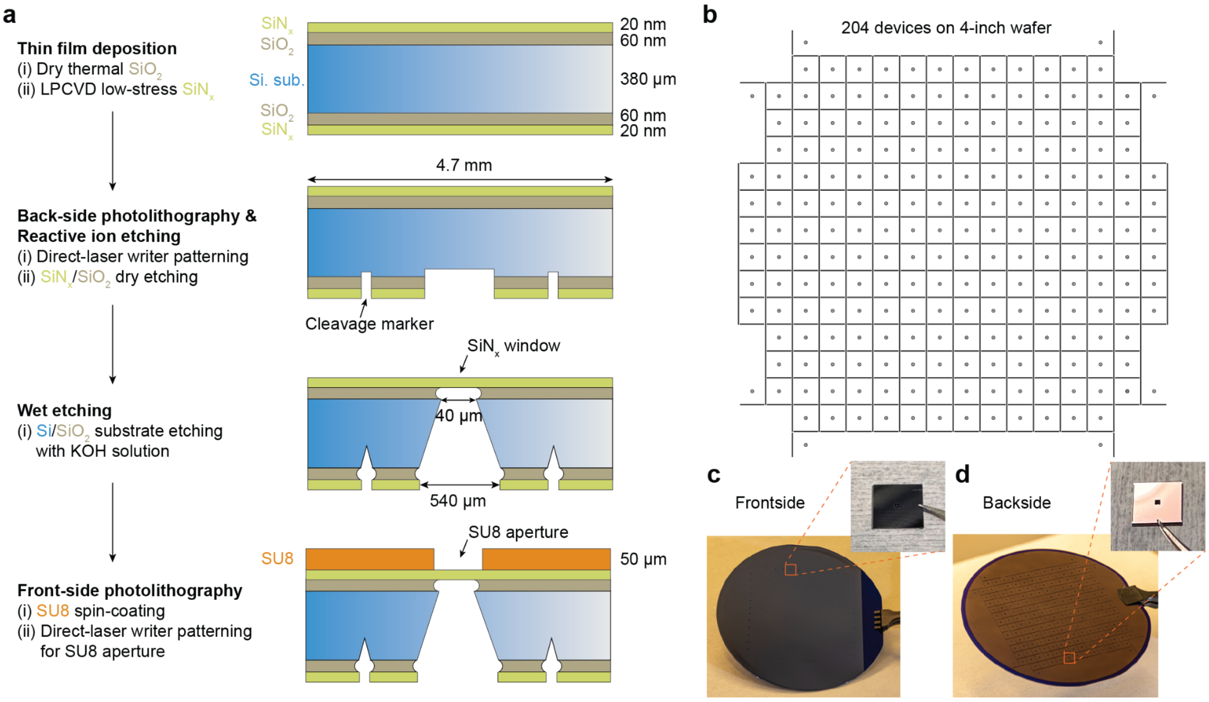

**Figure S1**. **Wafer-scale fabrication of low-noise devices**. **(a)** Microfabrication process flow to prepare suspending silicon nitride ($SiN_x$) windows using photolithography and reactive ion etching, followed by SU8 coating and aperture shaping in the SU8 layer via mask-less photolithography. **(b)** Designed pattern on the backside of the wafer. **(c, d)** Optical photos of the **(c)** frontside and **(d)** backside of SU8-coated wafer and a single chip (inset).

##### 1.2 Engineering of SU8 geometry with two-photon greyscale lithography

To further suppress capacitive noise originating from the electrolyte-accessible membrane area, we investigate non-cylindrical SU8 aperture geometries fabricated using two-photon polymerisation (TPP) lithography (Figure S2), benefitting from the compatibility of SU8 photoresist with the TPP method. While reducing the diameter of conventional cylindrical apertures decreases exposed $SiN_x$ membrane area, excessively small apertures can hinder electrolyte wetting and promote air bubble trapping, limiting practical device usability. To overcome this trade-off, TPP lithography was employed to fabricate tapered SU8 apertures with a 60 µm top opening and a 10 µm bottom opening (Figure S2.a), thereby preserving efficient liquid access while minimizing the membrane area exposed to the electrolyte. The fabrication process relies on localized two-photon-induced crosslinking of SU8 resist, followed by selective removal of uncured material during development. Electrical noise measurements demonstrate that devices incorporating tapered apertures exhibit reduced PSD and lower RMS noise compared with devices containing cylindrical apertures with same or even smaller top openings (Figure S2.d), further validating aperture geometry engineering as an effective strategy for minimizing chip capacitance and improving high-bandwidth recording performance.

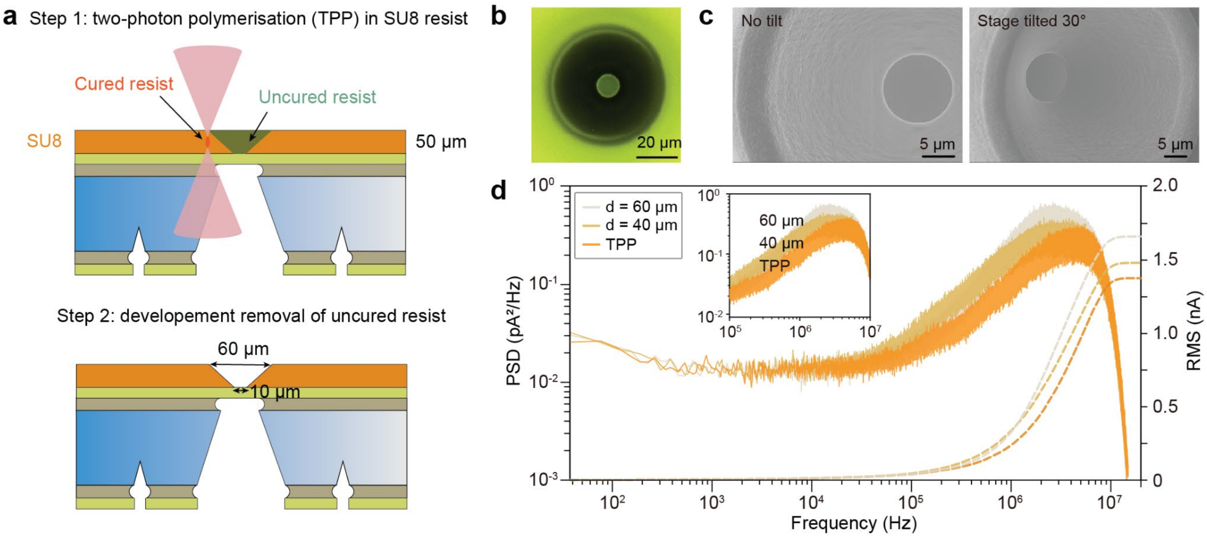

**Figure S2. Geometry engineering of SU8 aperture using the two-photon polymerisation (TPP) fabrication method. (a)** Process flow to expose uncured SU8 resist using two-photon lithography method and fabricate SU8 aperture in tapered shape, with top diameter of 60 µm and bottom diameter of 10 µm. **(b)** Optical image of the SU8-coated chip. **(c)** Scanning electron microscopy (SEM) images of the tapered SU8 aperture. **(d)** Noise power spectral density (PSD) and root-mean-square (RMS) noise comparison among the low-noise chips with TPP-shaped tapered SU8 aperture and cylindrical aperture that has different diameters.

##### 1.3 Noise comparison between various low-noise solid-state nanopore devices

We evaluate the effectiveness of the SU8-based low-noise chip by comparing its electrical noise characteristics with some representative solid-state nanopore architectures employing different capacitance reduction strategies, including silicone-passivated chips and commercial glass-supported nanopore devices, and compare their performances with the conventional bare $SiN_x$ chip (Figure S3). All devices have nanopores with diameters of 5–7 nm and are measured in 1 M KCl electrolyte (pH 6.0) using an Axopatch 200B amplifier operated at 100 kHz bandwidth and 200 kHz sampling frequency. While silicone coating and glass-supported designs reduce parasitic capacitance relative to bare $SiN_x$ membranes, the SU8-coated device achieves nearly the lowest baseline current noise and the noise power spectral density across the measured frequency range. Considering the much larger area of electrolyte-exposed $SiN_x$ membrane in the glass chip (Figure S3.c), the SU8 coating potentially presents better noise reduction effects compared with glass substrate. Unlike silicone-based passivation approaches that rely on manual coating on individual chips and suffer from uncontrollable thickness and coating area, as well as limited compatibility with high-bandwidth measurement systems, the SU8 approach enables wafer-scale, lithographically defined fabrication with improved device reproducibility. This comparison highlights SU8-based dielectric engineering as a scalable route toward simultaneously achieving low-noise performance and compatibility with ultra-high-bandwidth solid-state nanopore sensing.

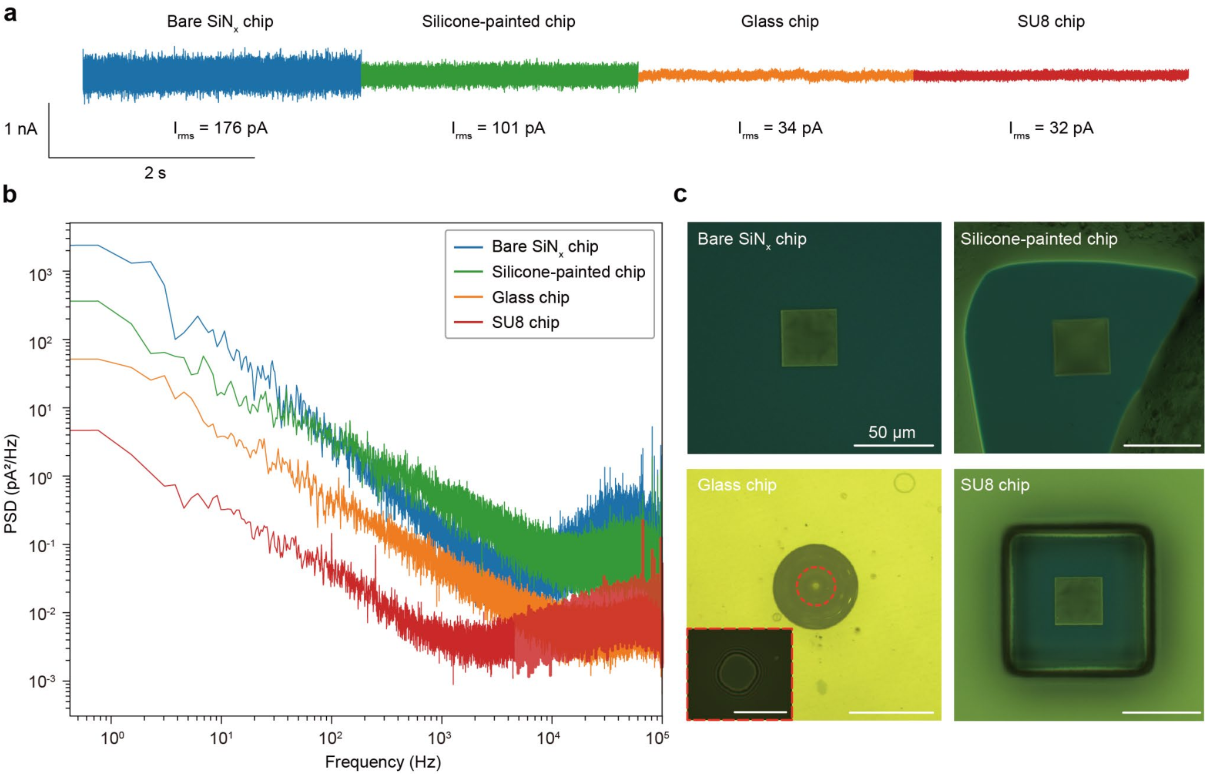

**Figure S3. Comparison of noise levels among various $SiN_x$ membrane-based nanopore devices. (a)** Representative ionic current trace and PSD from four types of nanopore devices, including the bare $SiN_x$ membrane chip, silicone-painted nanopore chip, commercial glass nanopore chip and SU8-coated $SiN_x$ nanopore chip. All chips have pore diameters between 5-7 nm and are tested with custom flowcell in 1M KCl solution (pH 6.0) using Axopatch 200B instrument with 100 kHz bandwidth and 200 kHz sampling frequency. Due to the uncontrollable silicone thickness, the painted nanopore chip is not compatible with the 10 MHz-bandwidth amplifier system. **(b)** Noise PSD comparison among four types of nanopore devices. **(c)** Optical images of the four types of nanopore devices. Inset of the glass chip is the higher magnification view of the suspending $SiN_x$ membrane where nanopore is fabricated. Scale bars in **(c)** are 50 μm.

##### 1.4 Wafer-scale mapping of SU8 thickness

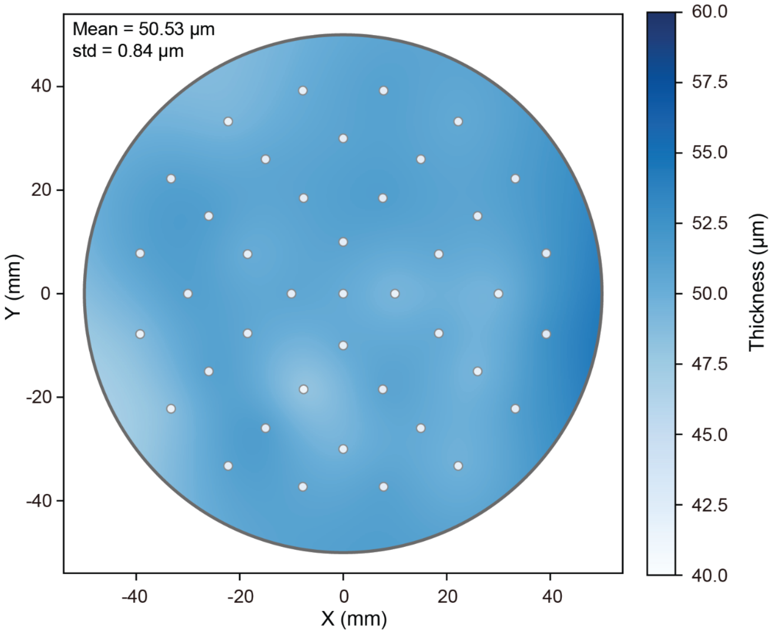

**Figure S4. Uniformity characterisation of the SU8 coating layer.** Wafer-scale mapping of SU8 thickness using Filmetrics F54 instrument and a 41-point measurement protocol is applied. Spin-coating parameters are set for a target thickness of 50 μm, and the measured average thickness across all points result in 50.53 $\pm$ 0.84 μm, indicating a good controllability and uniformity of SU8 coating.

##### 1.5 System capacitance measurement and calculation

The effective capacitance of the nanopore measurement system is quantified using a triangle-wave voltage stimulation protocol on the Elements amplifier and the current response is recorded (Figure S5). All measurements are performed in 1M KCl buffered solution (10 mM Tris-HCl and 1 mM EDTA, pH 8.0) under 250 kHz bandwidth. A symmetric triangular waveform of defined amplitude of 100 mV and 5 Hz sweeping frequency is applied across the nanopore, which leads to a constant voltage sweep rate (slope = 2000 mV/s). By modelling the nanopore system with an equivalent electrical circuit with a resistor $R$ and a capacitor $C$ in parallel, the current response of the system can be expressed as

$$I(t) = \frac{V(t)}{R} + C \cdot \frac{dV(t)}{dt}$$

where the first term stands for the resistive contribution and the second for capacitive contribution. Under the triangle-wave stimulus, the capacitive current response is expected to be constant and proportional to the voltage sweep rate $\alpha = \frac{dV(t)}{dt}$.

To extract the total capacitance of the system, the current response was segmented into rising and falling regions of the linearly swept voltage. The mean current values in each region were determined by excluding the transition regions where voltage polarity changes. The differences of mean current values $\Delta I$ were then used to compute the capacitance as $C = \frac{\Delta I}{2\alpha}$.

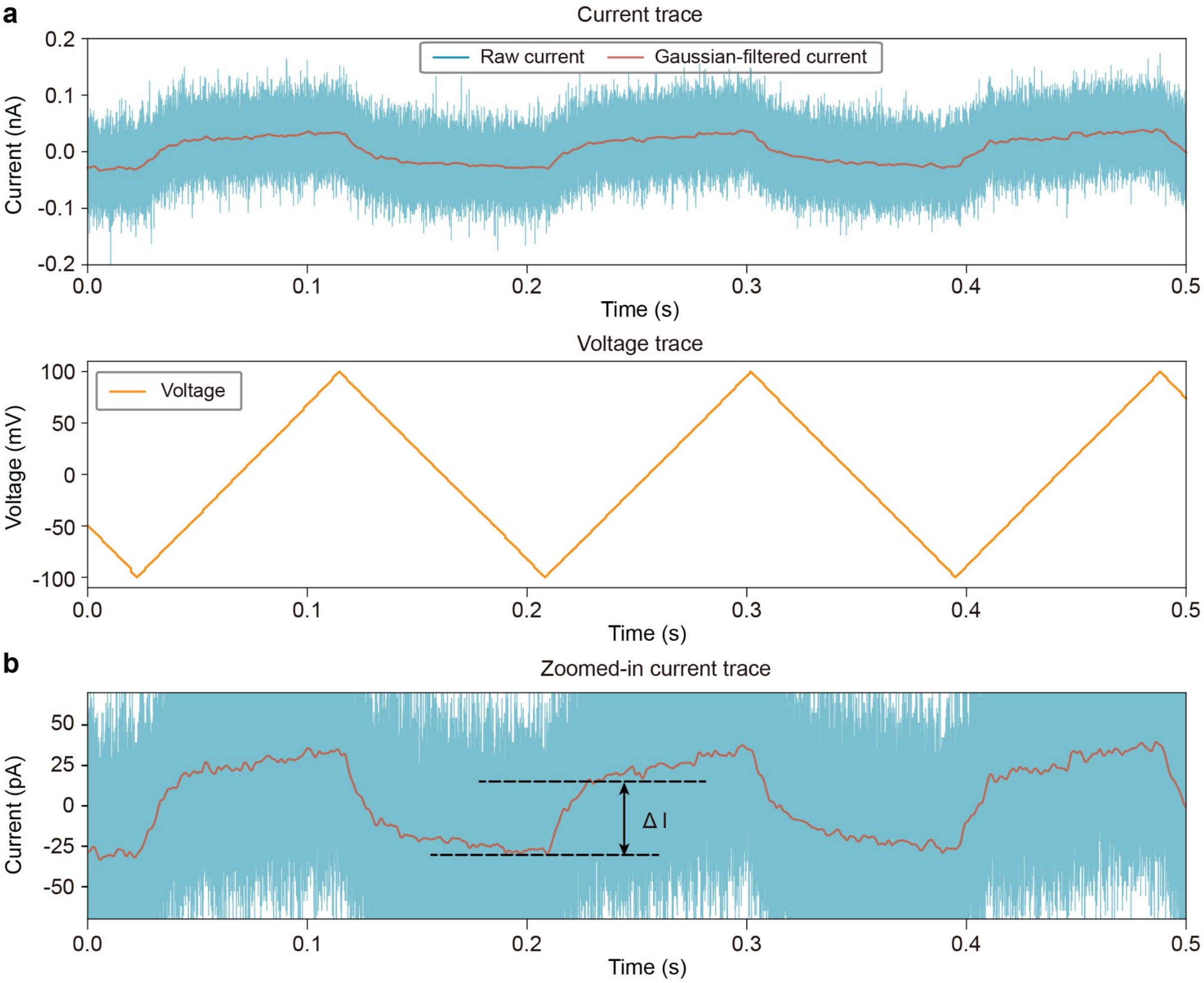

**Figure S5**. **Measurement of system capacitance using triangle-wave voltage protocol. (a)** Representative ionic current trace (top) of a low-noise chip $(d = 40\ \mu m,\ t = 50\ \mu m)$, measured in response to the triangle voltage wave (bottom). Raw current is collected under sampling frequency of 250 kHz via Elements instrument, overlaid with Gaussian-filtered data using sigma = 200. **(b)** Zoomed-in view of the ionic current trace and schematic representation of current change when voltage polarity reverses.

Besides direct measurement of the system capacitance, we also estimate the chip capacitance by considering the equivalent electrical model and geometrical factor of each individual layer composing the nanopore chip (Figure S6). The calculated chip capacitance exhibits the same dependency on the SU8 aperture size and coating thickness as the results obtained from triangle-wave measurement, with similar values from a few pF to hundreds of pF.

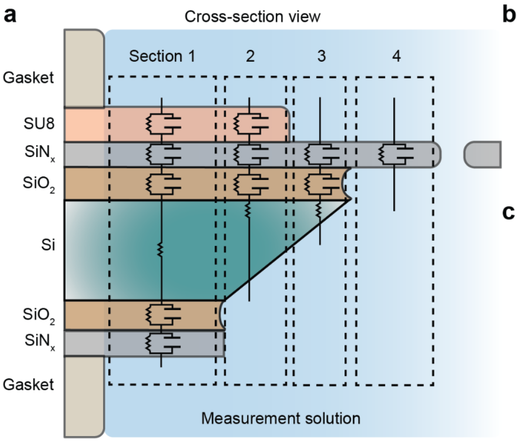

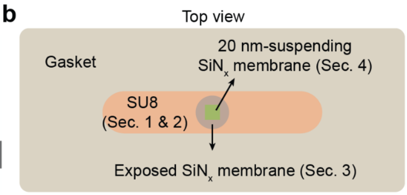

c

| Components | Relative Permittivity | Thickness |
|---|---|---|
| SU8 | 3.28 | 50 μm |
| Top layer $SiN_x$ | 7.5 | 20 nm |
| Top layer $SiO_2$ | 3.9 | 60 nm |
| Bottom layer $SiO_2$ | 3.9 | 60 nm |
| Bottom layer $SiN_x$ | 7.5 | 20 nm |

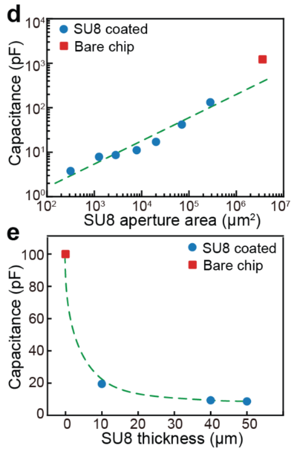

**Figure S6. Estimation of chip capacitance based on simplified equivalent electrical circuit model. (a)** Cross-sectional schematics of the electrical circuit model superimposed on the physical geometry of the chip. The plot is not in scale. **(b)** Top view of the electrolyte-exposed chip surface when assembled in flowcell and sandwiched by the gaskets. **(c)** Material properties of each individual component for capacitance calculation. **(d, e)** Calculated chip capacitance as a function of aperture diameter and SU8 thickness. The dashed lines are visual guides for the trend of capacitance change. Coating thickness is set at 50 μm when tuning the aperture size. Aperture size is 60 μm when varying the thickness.

#### Section 2. Design of deep learning framework to recover embedded transient feature in high-bandwidth nanopore translocation signal

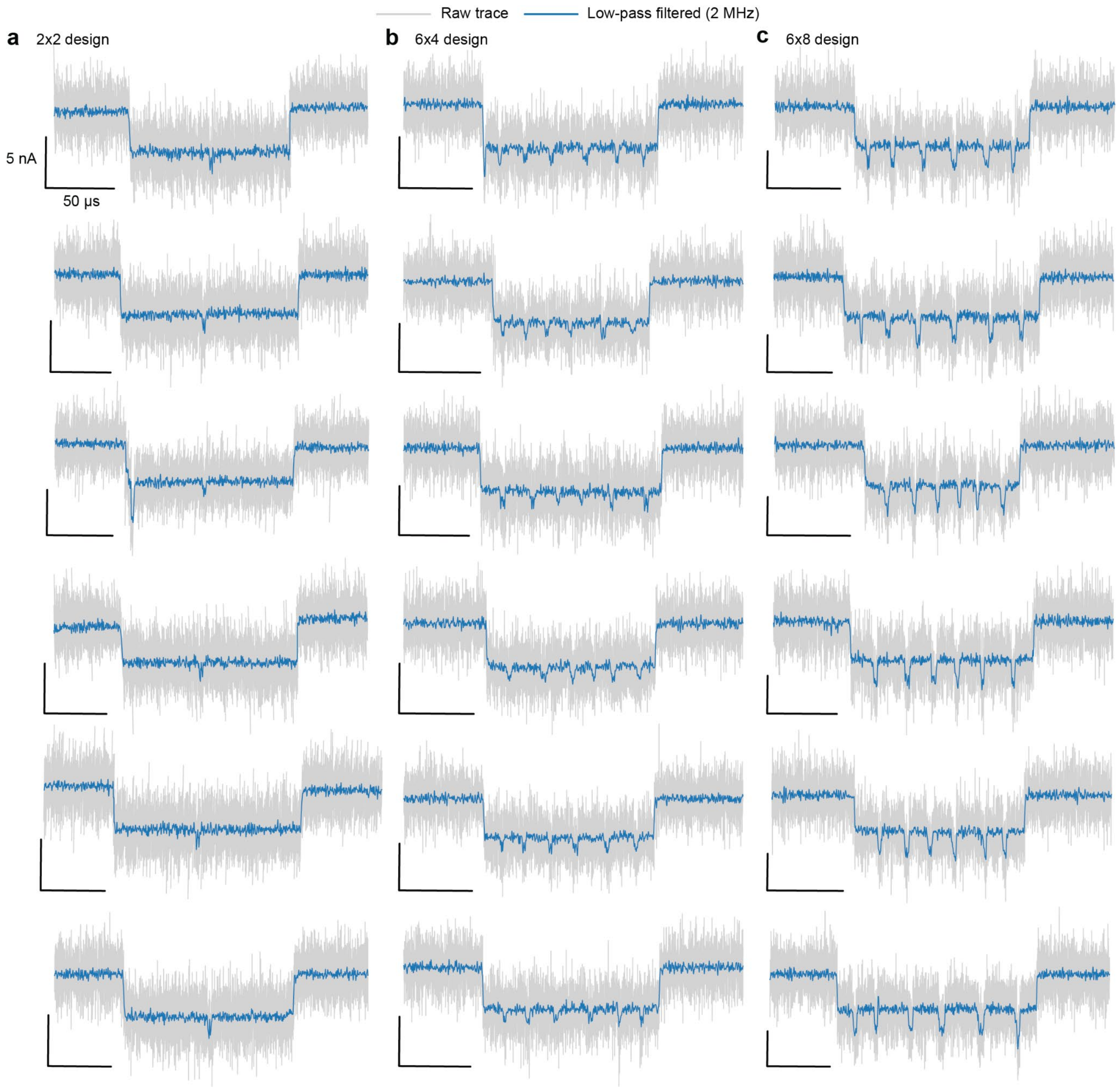

**Figure S7. Representative events of three types of DNA molecules carrying dumbbell nanostructures.** Translocation events of DNA constructs with **(a)** 2x2 design, **(b)** 6x4 design and **(c)** 6x8 design are measured under 40 MHz sampling rate (grey). Low-pass filtered current traces with cutoff frequency of 2 MHz are overlaid for better visuality. The raw trace records transient information with high temporal resolution, while the high-frequency noise embeds the information and makes it less distinguishable from the noise. Applying a low-pass filter enhances the signal-to-noise ratio but degrades the signal due to smoothing effect, making the transient features in signal indistinguishable.

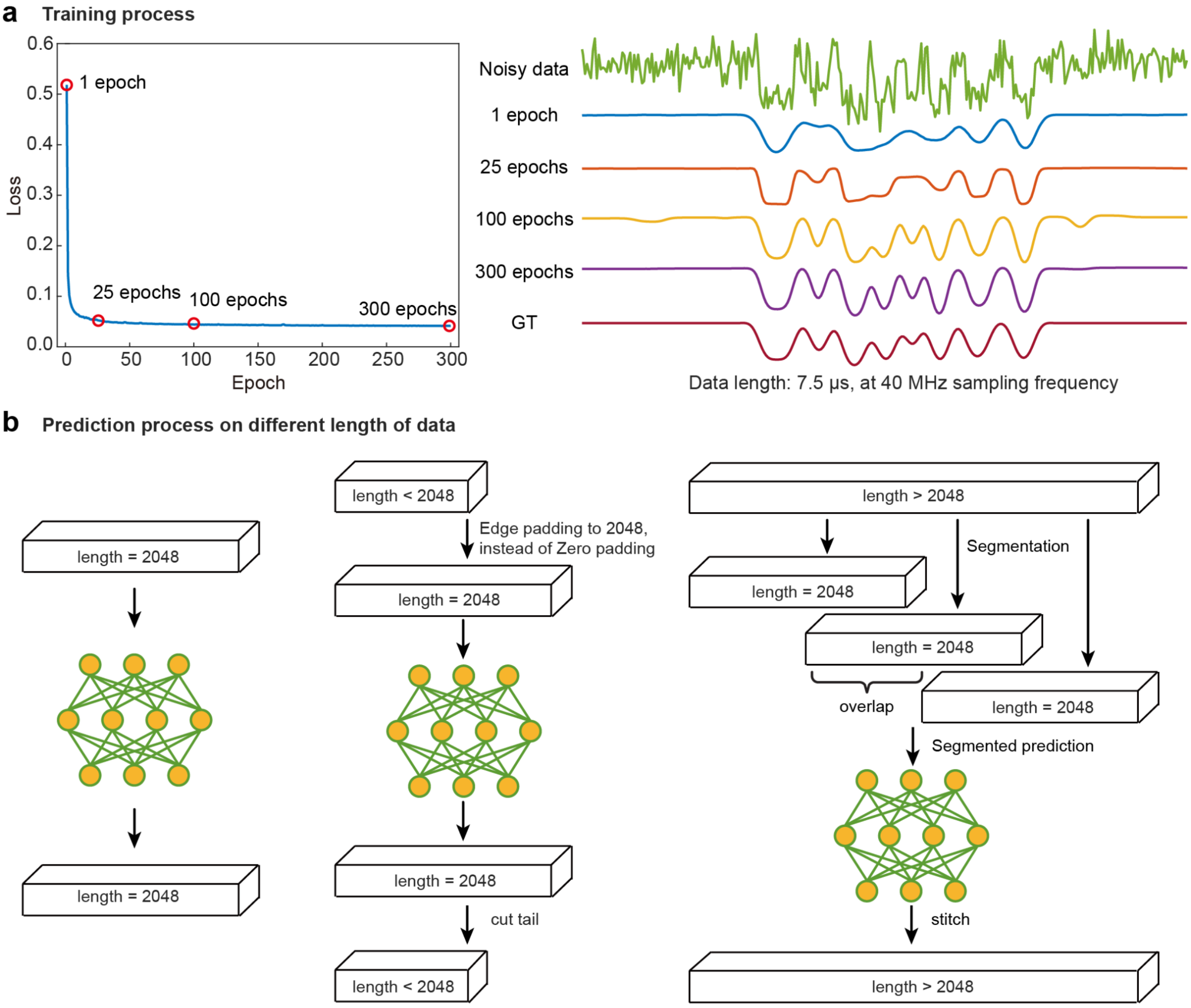

**Figure S8. Training and prediction phases of the deep learning model. (a)** Loss curve and reconstructed signal upon increasing epoch numbers. As the number of iterations increases, the network converges rapidly within a few dozen epochs, while the recovered high-frequency details gradually become richer and remain stable after 100 epochs. GT, Ground truth. **(b)** Schematics of the prediction process when different lengths of data are presented. The network reads 2048 data points at a time, so for data that matches this read length, the network can make predictions directly. For data shorter than this length, the network will extend the data to 2048 at the end by edge padding, and then crop it after the network makes a prediction. The edge padding is to extend the data length by copying the last point of the raw data, which is often used to avoid sharp edge transition when sampling near the borders of the zero padding. For data longer than this length, the network performs segmented predictions, and keeps a 30% overlap between each segment.

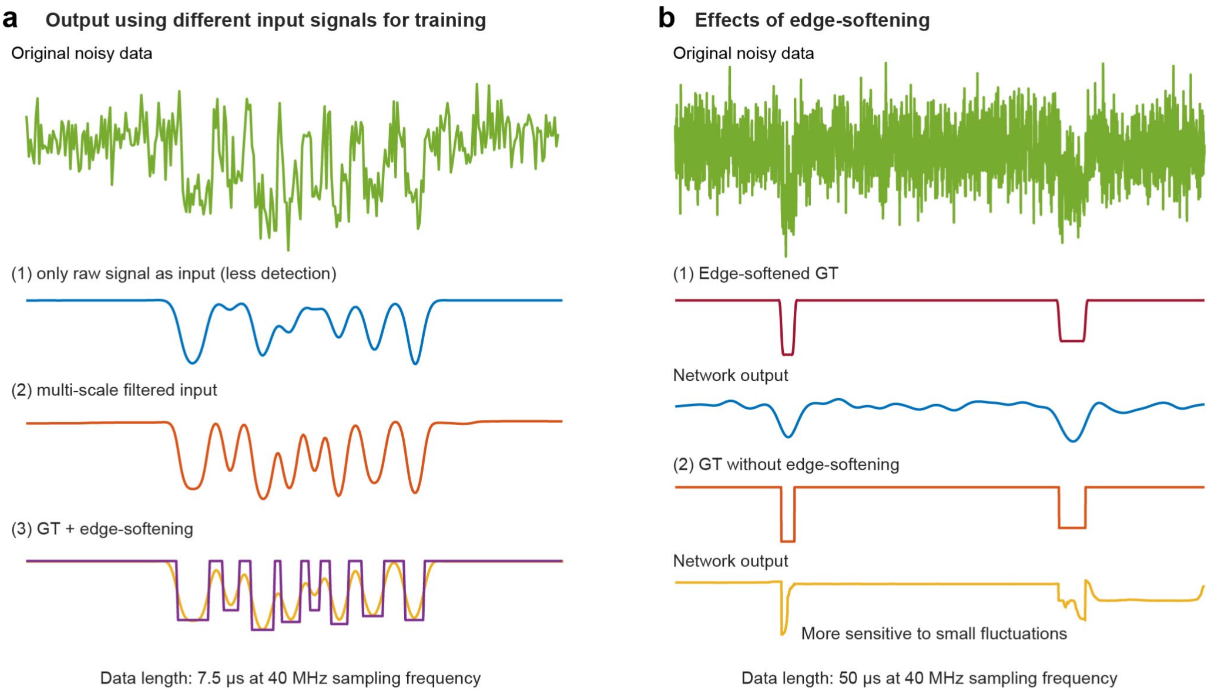

**Figure S9. Effect of data simulation and preparation methods on the reconstructed signal. (a)** Comparison of network training performance between multi-scale filtered input and single-channel raw data input.. The initialization parameters for the network training remain consistent. When training on single-channel raw data, the original channel was repeated multiple times to make it consistent with the multi-channel input. In terms of performance, the multi-channel input strategy expands the receptive field of the network by integrating information from different noise scales, thus exhibiting better sensitivity compared to the single-channel approach.**(b)** The effects of edge-softening on the GT for network performance. The sharp edges of square wave signals (GT) will make the network overfit the signal edge and become very sensitive to weak transitions, similar to the Ringing artifacts in frequency domain analysis. The softening can effectively alleviate this problem.

#### Section 3. Design and synthesis of DNA molecules carrying dumbbell nanostructures

##### 3.1 DNA sequence details of all strands

Three designs of nanostructured DNA molecules are studied in this work. The first benchmarking design (2x2 design) consists of 2 marker regions, with 2 dumbbells per marker region. The other two designs that are used to test and validate the codesigned platform contain 6 marker regions, with Design 1 consisting of 4 uniformly spaced dumbbells per marker region (6x4 design) and Design 2 consisting of 8 uniformly spaced dumbbells per marker region (6x8 design). As these two designs share the same marker regions, they therefore present the same marker window but different dumbbell-to-dumbbell distances inside the marker, with Design 1 having larger gaps than Design 2.

To synthesize all three designs of DNA molecules carrying dumbbell nanostructures, oligonucleotides that can form self-pairing hairpins are designed to replace the complementary staples. We adopted the 190 complementary oligonucleotide sequences and the sequences for benchmarking 2x2 design from previous work[1], and the sequence information is provided in **Table S1** and **Table S2**, respectively. For the two testing designs (**Table S3**,**S4**), we further modified the single dumbbell unit based on the benchmark design by adding two more C-G pairs to the self-pairing dumbbell backbone, as shown in Figure 3 and Figure 4 in main text. All designed DNA-dumbbells are evaluated using the NUPACK suite[2,3], to minimize secondary structure formation and optimize folding efficiency. This modification improves the thermal stability of DNA nanostructures, which will be further discussed in **Section 3.4**. In addition, with 20 bp center-to-center distance between neighbouring dumbbells, the new 6x8 design presents further reduced dumbbell-to-dumbbell gaps (4-8 nt gaps between the side-arms) compared with the molecules tested in the previous work[1,4].

**Complementary oligonucleotides**

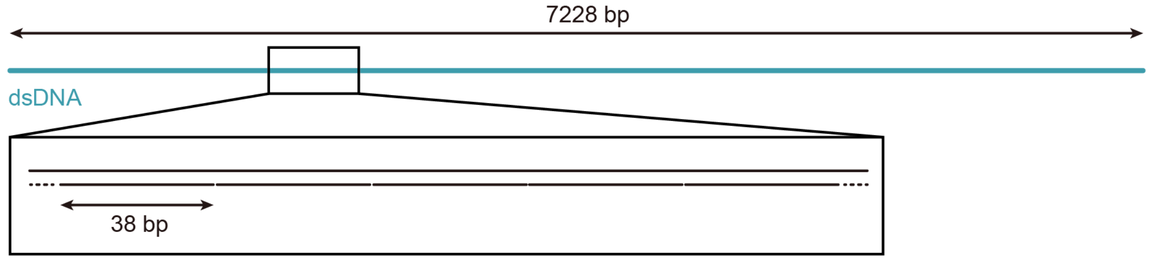

| Oligo. NO. | Sequence | Length (nt) | Oligo. NO. | Sequence | Length (nt) |
|---|---|---|---|---|---|
| 1 | TTTTCGTAATCATGGTCATAGCTGTTTCCTGTGTGAAATTGTTATC | 46 | 96 | CTTGAGCCATTTGGGAATTAGAGCCAGCAAAATCACCA | 38 |
| 2 | CGCTCACAATTCCACACAACATACGAGCCGGAAGCATA | 38 | 97 | GTAGCACCATTACCATTAGCAAGGCCGGAAACGTCACC | 38 |
| 3 | AAGTGTAAAGCCTGGGGTGCCTAATGAGTGAGCTAACT | 38 | 98 | AATGAAACCATCGATAGCAGCACCGTAATCAGTAGCGA | 38 |
| 4 | CACATTAATTGCGTTGCGCTCACTGCCCGCTTTCCAGT | 38 | 99 | CAGAATCAAGTTTGCCTTTAGCGTCAGACTGTAGCGCG | 38 |
| 5 | CGGGAAACCTGTCGTGCCAGCTGCATTAATGAATCGGC | 38 | 100 | TTTTCATCGGCATTTTCGGTCATAGCCCCCTTATTAGC | 38 |

| | | | | | |
|---|---|---|---|---|---|
| 6 | CAACGCGCGGGGAGAGGCGGTTTGCGTA TTGGGCGCCA | 38 | 101 | GTTTGCCATCTTTTCATAATCAAAATCAC CGGAACCAG | 38 |
| 7 | GGGTGGTTTTTCTTTTCACCAGTGAGACG GGCAACAGC | 38 | 102 | AGCCACCACCGGAACCGCCTCCCTCAG AGCCGCCACCC | 38 |
| 8 | TGATTGCCCTTCACCGCCTGGCCCTGAGA GAGTTGCAG | 38 | 103 | TCAGAACCGCCACCCTCAGAGCCACCA CCCTCAGAGCC | 38 |
| 9 | CAAGCGGTCCACGCTGGTTTGCCCCAGCA GGCGAAAAT | 38 | 104 | GCCACCAGAACCACCACCAGAGCCGCC GCCAGCATTGA | 38 |
| 10 | CCTGTTTGATGGTGGTTCCGAAATCGGCA AAATCCCTT | 38 | 105 | CAGGAGGTTGAGGCAGGTCAGACGATT GGCCTTGATAT | 38 |
| 11 | ATAAATCAAAAGAATAGCCCGAGATAGGG TTGAGTGTT | 38 | 106 | TCACAAACAAATAAATCCTCATTAAAGCC AGAATGGAA | 38 |
| 12 | GTTCCAGTTTGGAACAAGAGTCCACTATTA AAGAACGT | 38 | 107 | AGCGCAGTCTCTGAATTTACCGTTCCAG TAAGCGTCAT | 38 |
| 13 | GGACTCCAACGTCAAAGGGCGAAAAACCG TCTATCAGG | 38 | 108 | ACATGGCTTTTGATGATACAGGAGTGTA CTGGTAATAA | 38 |
| 14 | GCGATGGCCCACTACGTGAACCATCACCC AAATCAAGT | 38 | 109 | GTTTTAACGGGGTCAGTGCCTTGAGTAA CAGTGCCCGT | 38 |
| 15 | TTTTTGGGGTCGAGGTGCCGTAAAGCACT AAATCGGAA | 38 | 110 | ATAAACAGTTAATGCCCCCTGCCTATTTC GGAACCTAT | 38 |
| 16 | CCCTAAAGGGAGCCCCCGATTTAGAGCTT GACGGGGAA | 38 | 111 | TATTCTGAAACATGAAAGTATTAAGAGG CTGAGACTCC | 38 |
| 17 | AGCCGGCGAACGTGGCGAGAAAGGAAGG GAAGAAAGCG | 38 | 112 | TCAAGAGAAGGATTAGGATTAGCGGGGT TTTGCTCAGT | 38 |
| 18 | AAAGGAGCGGGCGCTAGGGCGCTGGCAA GTGTAGCGGT | 38 | 113 | ACCAGGCGGATAAGTGCCGTCGAGAGG GTTGATATAAG | 38 |
| 19 | CACGCTGCGCGTAACCACCACACCCGCC GCGCTTAATG | 38 | 114 | TATAGCCCGGAATAGGTGTATCACCGTA CTCAGGAGGT | 38 |
| 20 | CGCCGCTACAGGGCGCGTACTATGGTTGC TTTGACGAG | 38 | 115 | TTAGTACCGCCACCCTCAGAACCGCCAC CCTCAGAACC | 38 |
| 21 | CACGTATAACGTGCTTTCCTCGTTAGAATC AGAGCGGG | 38 | 116 | GCCACCCTCAGAGCCACCACCCTCATTT TCAGGGATAG | 38 |
| 22 | AGCTAAACAGGAGGCCGATTAAAGGGATT TTAGACAGG | 38 | 117 | CAAGCCCAATAGGAACCCATGTACCGTA ACACTGAGTT | 38 |
| 23 | AACGGTACGCCAGAATCCTGAGAAGTGTT TTTATAATC | 38 | 118 | TCGTCACCAGTACAAACTACAACGCCTG TAGCATTCCA | 38 |
| 24 | AGTGAGGCCACCGAGTAAAAGAGTCTGTC CATCACGCA | 38 | 119 | CAGACAGCCCTCATAGTTAGCGTAACGA TCTAAAGTTT | 38 |
| 25 | AATTAACCGTTGTAGCAATACTTCTTTGAT TAGTAATA | 38 | 120 | TGTCGTCTTTCCAGACGTTAGTAAATGA ATTTTCTGTA | 38 |
| 26 | ACATCACTTGCCTGAGTAGAAGAACTCAAA CTATCGGC | 38 | 121 | TGGGATTTTGCTAAACAACTTTCAACAGT TTCAGCGGA | 38 |
| 27 | CT/TGCTGGTAATATCCAGAACA/ATATTAC CGCCAGCCA | 40 | 122 | GTGAGAATAGAAAGGAACAACTAAAGGA ATTGCGAATA | 38 |
| 28 | TTGCAACAGGAAAAACGCTCATGGAAATA CCTACATTT | 38 | 123 | ATAATTTTTTCACGTTGAAAATCTCCAAA AAAAAGGCT | 38 |
| 29 | TGACGCTCAATCGTCTGAAATGGATTATTT ACATTGGC | 38 | 124 | CCAAAAGGAGCCTTTAATTGTATCGGTT TATCAGCTTG | 38 |
| 30 | AGATTCACCAGTCACACGACCAGTAATAAA AGGGACAT | 38 | 125 | CTTTCGAGGTGAATTTCTTAAACAGCTTG ATACCGATA | 38 |
| 31 | TCTGGCCAACAGAGATAGAACCCTTCTGA CCTGAAAGC | 38 | 126 | GTTGCGCCGACAATGACAACAACCATCG CCCACGCATA | 38 |
| 32 | GTAAGAATACGTGGCACAGACAATATTTTT GAATGGCT | 38 | 127 | ACCGATATATTCGGTCGCTGAGGCTTGC AGGGAGTTAA | 38 |
| 33 | ATTAGTCTTTAATGCGCGAACTGATAGCCC TAAAACAT | 38 | 128 | AGGCCGCTTTTGCGGGATCGTCACCCT CAGCAGCGAAA | 38 |
| 34 | CGCCATTAAAAATACCGAACGAACCACCA GCAGAAGAT | 38 | 129 | GACAGCATCGGAACGAGGGTAGCAACG GCTACAGAGGC | 38 |
| 35 | AAAACAGAGGTGAGGCGGTCAGTATTAAC ACCGCCTGC | 38 | 130 | TTTGAGGACTAAAGACTTTTTCATGAGG AAGTTTCCAT | 38 |
| 36 | AACAGTGCCACGCTGAGAGCCAGCAGCAA ATGAAAAAT | 38 | 131 | TAAACGGGTAAAATACGTAATGCCACTA CGAAGGCACC | 38 |
| 37 | CTAAAGCATCACCTTGCTGAACCTCAAATA TCAAACCC | 38 | 132 | AACCTAAAACGAAAGAGGCAAAAGAATA CACTAAAACA | 38 |

| | | | | | |
|---|---|---|---|---|---|
| 38 | TCAATCAATATCTGGTCAGTTGGCAAATCA ACAGTTGA | 38 | 133 | CTCATCTTTGACCCCCAGCGATTATACC AAGCGCGAAA | 38 |
| 39 | AAGGAATTGAGGAAGGTTATCTAAAATATC TTTAGGAG | 38 | 134 | CAAAGTACAACGGAGATTTGTATCATCG CCTGATAAAT | 38 |
| 40 | CACTAACAACTAATAGATTAGAGCCGTCAA TAGATAAT | 38 | 135 | TGTGTCGAAATCCGCGACCTGCTCCATG TTACTTAGCC | 38 |
| 41 | ACATTTGAGGATTTAGAAGTATTAGACTTT ACAAACAA | 38 | 136 | GGAACGAGGCGCAGACGGTCAATCATA AGGGAACCGAA | 38 |
| 42 | TTCGACAACTCGTATTAAATCCTTTGCCCG AACGTTAT | 38 | 137 | CTGACCAACTTTGAAAGAGGACAGATGA ACGGTGTACA | 38 |
| 43 | TAATTTTAAAAGTTTGAGTAACATTATCATT TTGCGGA | 38 | 138 | GACCAGGCGCATAGGCTGGCTGACCTT CATCAAGAGTA | 38 |
| 44 | ACAAAGAAACCACCAGAAGGAGCGGAATT ATCATCATA | 38 | 139 | ATCTTGACAAGAACCGGATATTCATTAC CCAAATCAAC | 38 |
| 45 | TTCCTGATTATCAGATGATGGCAATTCATC AATATAAT | 38 | 140 | GTAACAAAGCTGCTCATTCAGTGAATAA GGCTTGCCCT | 38 |
| 46 | CCTGATTGTTTGGATTATACTTCTGAATAA TGGAAGGG | 38 | 141 | GACGAGAAACACCAGAACGAGTAGTAAA TTGGGCTTGA | 38 |
| 47 | TTAGAACCTACCATATCAAAATTATTTGCA CGTAAAAC | 38 | 142 | GATGGTTTAATTTCAACTTTAATCATTGT GAATTACCT | 38 |
| 48 | AGAAATAAAGAAATTGCGTAGATTTTCAGG TTTAACGT | 38 | 143 | TATGCGATTTTAAGAACTGGCTCATTATA CCAGTCAGG | 38 |
| 49 | CAGATGAATATACAGTAACAGTACCTTTTA CATCGGGA | 38 | 144 | ACGTTGGGAAGAAAAATCTACGTTAATA AAACGAACTA | 38 |
| 50 | GAAACAATAACGGATTCGCCTGATTGCTTT GAATACCA | 38 | 145 | ACGGAACAACATTATTACAGGTAGAAAG ATTCATCAGT | 38 |
| 51 | AGTTACAAAATCGCGCAGAGGCGAATTAT TCATTTCAA | 38 | 146 | TGAGATTTAGGAATACCACATTCAACTAA TGCAGATAC | 38 |
| 52 | TTACCTGAGCAAAAGAAGATGATGAAACAA ACATCAAG | 38 | 147 | ATAACGCCAAAAGGAATTACGAGGCATA GTAAGAGCAA | 38 |
| 53 | AAAACAAAATTAATTACATTTAACAATTTCA TTTGAAT | 38 | 148 | CACTATCATAACCCTCGTTTACCAGACG ACGATAAAAA | 38 |
| 54 | TACCTTTTTTAATGGAAACAGTACATAAAT CAATATAT | 38 | 149 | CCAAAATAGCGAGAGGCTTTTGCAAAAG AAGTTTTGCC | 38 |
| 55 | GTGAGTGAATAACCTTGCTTCTGTAAATCG TCGCTATT | 38 | 150 | AGAGGGGGTAATAGTAAAATGTTTAGAC TGGATAGCGT | 38 |
| 56 | AATTAATTTTCCCTTAGAATCCTTGAAAACA TAGCGAT | 38 | 151 | CCAATACTGCGGAATCGTCATAAATATT CATTGAATCC | 38 |
| 57 | AGCTTAGATTAAGACGCTGAGAAGAGTCA ATAGTGAAT | 38 | 152 | CCCTCAAATGCTTTAAACAGTTCAGAAAA CGAGAATGA | 38 |
| 58 | TTATCAAAATCATAGGTCTGAGAGACTACC TTTTTAAC | 38 | 153 | CCATAAATCAAAAATCAGGTCTTTACCCT GACTATTAT | 38 |
| 59 | CTCCGGCTTAGGTTGGGTTATATAACTATA TGTAAATG | 38 | 154 | AGTCAGAAGCAAAGCGGATTGCATCAAA AAGATTAAGA | 38 |
| 60 | CTGATGCAAATCCAATCGCAAGACAAAGA ACGCGAGAA | 38 | 155 | GGAAGCCCGAAAGACTTCAAATATCGCG TTTTAATTCG | 38 |
| 61 | AACTTTTTCAAATATATTTTAGTTAATTTCA TCTTCTG | 38 | 156 | AGCTTCAAAGCGAACCAGACCGGAAGC AAACTCCAACA | 38 |
| 62 | ACCTAAATTTAATGGTTTGAAATACCGACC GTGTGATA | 38 | 157 | GGTCAGGATTAGAGAGTACCTTTAATTG CTCCTTTTGA | 38 |
| 63 | AATAAGGCGTTAAATAAGAATAAACACCGG AATCATAA | 38 | 158 | TAAGAGGTCATTTTTGCGGATGGCTTAG AGCTTAATTG | 38 |
| 64 | TTACTAGAAAAAGCCTGTTTAGTATCATAT GCGTTATA | 38 | 159 | CTGAATATAATGCTGTAGCTCAACATGTT TTAAATATG | 38 |
| 65 | CAAATTCTTACCAGTATAAAGCCAACGCTC AACAGTAG | 38 | 160 | CAACTAAAGTACGGTGTCTGGAAGTTTC ATTCCATATA | 38 |
| 66 | GGCTTAATTGAGAATCGCCATATTTAACAA CGCCAACA | 38 | 161 | ACAGTTGATTCCCAATTCTGCGAACGAG TAGATTTAGT | 38 |
| 67 | TGTAATTTAGGCAGAGGCATTTTCGAGCC AGTAATAAG | 38 | 162 | TTGACCATTAGATACATTTCGCAAATGGT CAATAACCT | 38 |
| 68 | AGAATATAAAGTACCGACAAAAGGTAAAGT AATTCTGT | 38 | 163 | GTTTAGCTATATTTTCATTTGGGGCGCG AGCTGAAAAG | 38 |
| 69 | CCAGACGACGACAATAAACAACATGTTCA GCTAATGCA | 38 | 164 | GTGGCATCAATTCTACTAATAGTAGTAG CATTAACATC | 38 |

| | | | | | |
|---|---|---|---|---|---|
| 70 | GAACGCGCCTGTTTATCAACAATAGATAAG TCCTGAAC | 38 | 165 | CAATAAATCATACAGGCAAGGCAAAGAA TTAGCAAAAT | 38 |
| 71 | AAGAAAAATAATATCCCATCCTAATTTACG AGCATGTA | 38 | 166 | TAAGCAATAAAGCCTCAGAGCATAAAGC TAAATCGGTT | 38 |
| 72 | GAAACCAATCAATAATCGGCTGTCTTTCCT TATCATTC | 38 | 167 | GTACCAAAAACATTATGACCCTGTAATAC TTTTGCGGG | 38 |
| 73 | CAAGAACGGGTATTAAACCAAGTACCGCA CTCATCGAG | 38 | 168 | AGAAGCCTTTATTTCAACGCAAGGATAA AAATTTTTAG | 38 |
| 74 | AACAAGCAAGCCGTTTTTATTTTCATCGTA GGAATCAT | 38 | 169 | AACCCTCATATATTTTAAATGCAATGCCT GAGTAATGT | 38 |
| 75 | TACCGCGCCCAATAGCAAGCAAATCAGAT ATAGAAGGC | 38 | 170 | GTAGGTAAAGATTCAAAAGGGTGAGAAA GGCCGGAGAC | 38 |
| 76 | TTATCCGGTATTCTAAGAACGCGAGGCGT TTTAGCGAA | 38 | 171 | AGTCAAATCACCATCAATATGATATTCAA CCGTTCTAG | 38 |
| 77 | CCTCCCGACTTGCGGGAGGTTTTGAAGCC TTAAATCAA | 38 | 172 | CTGATAAATTAATGCCGGAGAGGGTAGC TATTTTTGAG | 38 |
| 78 | GATTAGTTGCTATTTTGCACCCAGCTACAA TTTTATCC | 38 | 173 | AGATCTACAAAGGCTATCAGGTCATTGC CTGAGAGTCT | 38 |
| 79 | TGAATCTTACCAACGCTAACGAGCGTCTTT CCAGAGCC | 38 | 174 | GGAGCAAACAAGAGAATCGATGAACGG TAATCGTAAAA | 38 |
| 80 | TAATTTGCCAGTTACAAAATAAACAGCCAT ATTATTTA | 38 | 175 | CTAGCATGTCAATCATATGTACCCCGGT TGATAATCAG | 38 |
| 81 | TCCCAATCCAAATAAGAAACGATTTTTTGT TTAACGTC | 38 | 176 | AAAAGCCCCAAAAACAGGAAGATTGTAT AAGCAAATAT | 38 |
| 82 | AAAAATGAAAATAGCAGCCTTTACAGAGAG AATAACAT | 38 | 177 | TTAAATTGTAAACGTTAATATTTTGTTAAA ATTCGCAT | 38 |
| 83 | AAAAACAGGGAAGCGCATTAGACGGGAGA ATTAACTGA | 38 | 178 | TAAATTTTTGTTAAATCAGCTCATTTTTTA ACCAATAG | 38 |
| 84 | ACACCCTGAACAAAGTCAGAGGGTAATTG AGCGCTAAT | 38 | 179 | GAACGCCATCAAAAATAATTCGCGTCTG GCCTTCCTGT | 38 |
| 85 | ATCAGAGAGATAACCCACAAGAATTGAGTT AAGCCCAA | 38 | 180 | AGCCAGCTTTCATCAACATTAAATGTGA GCGAGTAACA | 38 |
| 86 | TAATAAGAGCAAGAAACAATGAAATAGCAA TAGCTATC | 38 | 181 | ACCCGTCGGATTCTCCGTGGGAACAAAC GGCGGATTGA | 38 |
| 87 | TTACCGAAGCCCTTTTTAAGAAAAGTAAGC AGATAGCC | 38 | 182 | CCGTAATGGGATAGGTCACGTTGGTGTA GATGGGCGCA | 38 |
| 88 | GAACAAAGTTACCAGAAGGAAACCGAGGA AACGCAATA | 38 | 183 | TCGTAACCGTGCATCTGCCAGTTTGAGG GGACGACGAC | 38 |
| 89 | ATAACGGAATACCCAAAAGAACTGGCATG ATTAAGACT | 38 | 184 | AGTATCGGCCTCAGGAAGATCGCACTCC AGCCAGCTTT | 38 |
| 90 | CCTTATTACGCAGTATGTTAGCAAACGTAG AAAATACA | 38 | 185 | CCGGCACCGCTTCTGGTGCCGGAAACC AGGCAAAGCGC | 38 |
| 91 | TACATAAAGGTGGCAACATATAAAAGAAAC GCAAAGAC | 38 | 186 | CATTCGCCATTCAGGCTGCGCAACTGTT GGGAAGGGCG | 38 |
| 92 | ACCACGGAATAAGTTTATTTTGTCACAATC AATAGAAA | 38 | 187 | ATCGGTGCGGGCCTCTTCGCTATTACGC CAGCTGGCGA | 38 |
| 93 | ATTCATATGGTTTACCAGCGCCAAAGACAA AAGGGCGA | 38 | 188 | AAGGGGGATGTGCTGCAAGGCGATTAA GTTGGGTAACG | 38 |
| 94 | CATTCAACCGATTGAGGGAGGGAAGGTAA ATATTGACG | 38 | 189 | CCAGGGTTTTCCCAGTCACGACGTTGTA AAACGACGGC | 38 |
| 95 | GAAATTATTCATTAAAGGTGAATTATCACC GTCACCGA | 38 | 190 | CAGTGCCAAGCTTGCATGCCTGCAGGT CGACTCTAGAGGATCTTTT | 46 |

**Table S1**. **Sequences of 190 complementary oligonucleotides for the M13mp18 single-strand scaffold**.

**2x2 design**

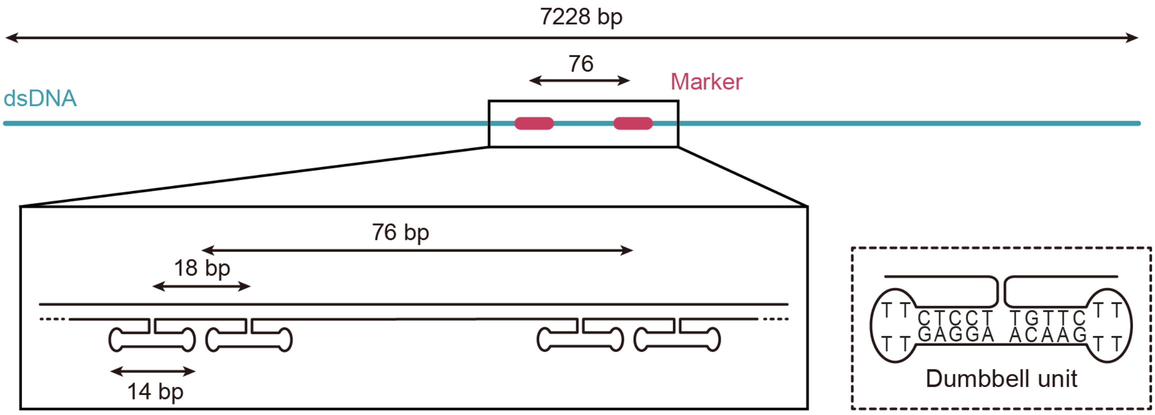

| Dumbbell number | Sequence | Length (nt) | Oligo positions to be replaced |
|---|---|---|---|
| 1 | GAAATTATTCTCCTCTTTTGAGGAACAAGTTTTCTTGTATTAAAGGT | 47 | 95 |
| 2 | GAATTATCATCCTCTTTTGAGGAACAAGTTTTCTTGTCCGTCACCGA | 47 | |
| 3 | GTAGCACCATCCTCTTTTGAGGAACAAGTTTTCTTGTTTACCATTAG | 47 | 97 |
| 4 | CAAGGCCGGTCCTCTTTTGAGGAACAAGTTTTCTTGTAAACGTCACC | 47 | |

**Table S2**. **Sequences of the oligonucleotides to form 2x2 design with in total 4 dumbbells**. In each marker region, the original complementary oligonucleotides at corresponding positions from Table S1 are replaced with the oligonucleotides here, which could form dumbbell hairpin structures attached to the dsDNA backbone after annealing.

**6x4 design**

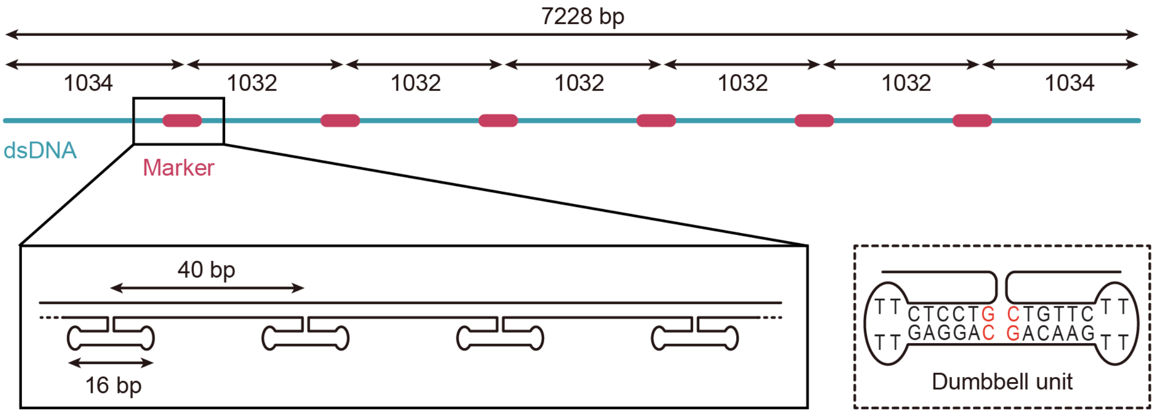

| Marker NO. | Sequence | Length (nt) | Oligo position to be replaced |
|---|---|---|---|
| 1 | ACATCACTTGGTCCTCTTTTGAGGACGACAAGTTTTCTTGTCCCTGAGTAGA | 52 | 26-30 |

| | | | |
|---|---|---|---|
| | AGAACTCAAACTATCGGCCT | 20 | |
| | TGCTGGTAATGTCCTCTTTTGAGGACGACAAGTTTTCTTGTCATCCAGAACA | 52 | |
| | ATATTACCGCCAGCCATTGC | 20 | |
| | AACAGGAAAAGTCCTCTTTTGAGGACGACAAGTTTTCTTGTCACGCTCATGG | 52 | |
| | AAATACCTACATTTTGACGC | 20 | |
| | TCAATCGTCTGTCCTCTTTTGAGGACGACAAGTTTTCTTGTCGAAATGGATT | 52 | |
| | ATTTACATTGGCAGATTCAC | 20 | |
| | CAGTCACACGACCAGTAATAAAAGGGACAT | 30 | |
| 2 | TTACCTGAGCAAAAGAAGATGATGAAACAAACATCAAGAAAACA | 44 | 52-57 |
| | AAATTAATTAGTCCTCTTTTGAGGACGACAAGTTTTCTTGTCCATTTAACAA | 52 | |
| | TTTCATTTGAATTACCTTTT | 20 | |
| | TTAATGGAAAGTCCTCTTTTGAGGACGACAAGTTTTCTTGTCCAGTACATAA | 52 | |
| | ATCAATATATGTGAGTGAAT | 20 | |
| | AACCTTGCTTGTCCTCTTTTGAGGACGACAAGTTTTCTTGTCCTGTAAATCG | 52 | |
| | TCGCTATTAATTAATTTTCC | 20 | |
| | CTTAGAATCCGTCCTCTTTTGAGGACGACAAGTTTTCTTGTCTTGAAAACAT | 52 | |
| | AGCGATAGCTTAGATTAAGA | 20 | |
| | CGCTGAGAAGAGTCAATAGTGAAT | 24 | |
| 3 | TGAATCTTACCAACGCTAACGAGCGTCTTTCCAGAGCCTAATTTGCCAGT | 50 | 79-85 |
| | TACAAAATAAGTCCTCTTTTGAGGACGACAAGTTTTCTTGTCACAGCCATAT | 52 | |
| | TATTTATCCCAATCCAAATA | 20 | |
| | AGAAACGATTGTCCTCTTTTGAGGACGACAAGTTTTCTTGTCTTTTGTTTAA | 52 | |
| | CGTCAAAAATGAAAATAGCA | 20 | |
| | GCCTTTACAGGTCCTCTTTTGAGGACGACAAGTTTTCTTGTCAGAGAATAAC | 52 | |
| | ATAAAAACAGGGAAGCGCAT | 20 | |
| | TAGACGGGAGGTCCTCTTTTGAGGACGACAAGTTTTCTTGTCAATTAACTGA | 52 | |
| | ACACCCTGAACAAAGTCAGA | 20 | |

| | | | |
|---|---|---|---|
| | GGGTAATTGAGCGCTAATATCAGAGAGATAACCCACAAGAATTGAGTTAAGCCCAA | 56 | |
| 4 | TCACAAACAAATAAATCCTCATTAAAGCCAGAATGGAAAGCGCAGTCTCTGAATTT | 56 | 106-112 |
| | ACCGTTCCAGGTCCTCTTTTGAGGACGACAAGTTTTCTTGTCTAAGCGTCAT | 52 | |
| | ACATGGCTTTTGATGATACA | 20 | |
| | GGAGTGTACTGTCCTCTTTTGAGGACGACAAGTTTTCTTGTCGGTAATAAGT | 52 | |
| | TTTAACGGGGTCAGTGCCTT | 20 | |
| | GAGTAACAGTGTCCTCTTTTGAGGACGACAAGTTTTCTTGTCGCCCGTATAA | 52 | |
| | ACAGTTAATGCCCCCTGCCT | 20 | |
| | ATTTCGGAACGTCCTCTTTTGAGGACGACAAGTTTTCTTGTCCTATTATTCT | 52 | |
| | GAAACATGAAAGTATTAAGA | 20 | |
| | GGCTGAGACTCCTCAAGAGAAGGATTAGGATTAGCGGGGTTTTGCTCAGT | 50 | |
| 5 | CAAAGTACAACGGAGATTTGTATC | 24 | 134-139 |
| | ATCGCCTGATGTCCTCTTTTGAGGACGACAAGTTTTCTTGTCAAATTGTGTC | 52 | |
| | GAAATCCGCGACCTGCTCCA | 20 | |
| | TGTTACTTAGGTCCTCTTTTGAGGACGACAAGTTTTCTTGTCCCGGAACGAG | 52 | |
| | GCGCAGACGGTCAATCATAA | 20 | |
| | GGGAACCGAAGTCCTCTTTTGAGGACGACAAGTTTTCTTGTCCTGACCAACT | 52 | |
| | TTGAAAGAGGACAGATGAAC | 20 | |
| | GGTGTACAGAGTCCTCTTTTGAGGACGACAAGTTTTCTTGTCCCAGGCGCAT | 52 | |
| | AGGCTGGCTGACCTTCATCA | 20 | |
| | AGAGTAATCTTGACAAGAACCGGATATTCATTACCCAAATCAAC | 44 | |
| 6 | ACAGTTGATTCCCAATTCTGCGAACGAGTA | 30 | 161-165 |
| | GATTTAGTTTGTCCTCTTTTGAGGACGACAAGTTTTCTTGTCGACCATTAGA | 52 | |
| | TACATTTCGCAAATGGTCAA | 20 | |
| | TAACCTGTTTGTCCTCTTTTGAGGACGACAAGTTTTCTTGTCAGCTATATTT | 52 | |
| | TCATTTGGGGCGCGAGCTGA | 20 | |
| | AAAGGTGGCAGTCCTCTTTTGAGGACGACAAGTTTTCTTGTCTCAATTCTAC | 52 | |

| | | |
|---|---|---|
| | TAATAGTAGTAGCATTAACA | 20 |
| | TCCAATAAATGTCCTCTTTTGAGGACGACAAGTTTTCTTGTCCATACAGGCA | 52 |
| | AGGCAAAGAATTAGCAAAAT | 20 |

**Table S3**. **Sequences of the oligonucleotides to form 6x4 design (design 1) with 6 marker regions and 4 dumbbells per marker region**.

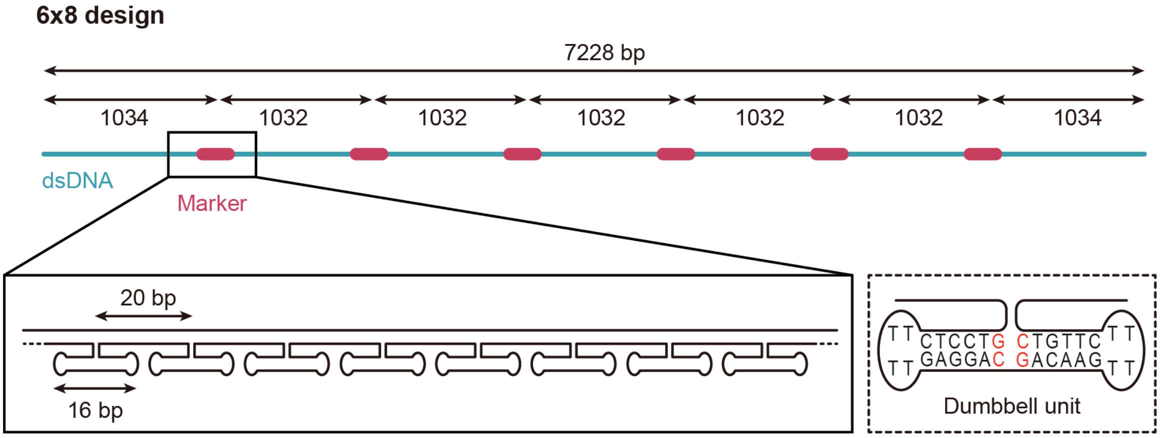

| Marker NO. | Sequence | Length (nt) | Oligo position to be replaced |
|---|---|---|---|
| 1 | ACATCACTTGGTCCTCTTTTGAGGACGACAAGTTTTCTTGTCCCTGAGTAGA | 52 | 26-30 |
| | AGAACTCAAAGTCCTCTTTTGAGGACGACAAGTTTTCTTGTCCTATCGGCCT | 52 | |
| | TGCTGGTAATGTCCTCTTTTGAGGACGACAAGTTTTCTTGTCATCCAGAACA | 52 | |
| | ATATTACCGCGTCCTCTTTTGAGGACGACAAGTTTTCTTGTCCAGCCATTGC | 52 | |
| | AACAGGAAAAGTCCTCTTTTGAGGACGACAAGTTTTCTTGTCACGCTCATGG | 52 | |
| | AAATACCTACGTCCTCTTTTGAGGACGACAAGTTTTCTTGTCATTTTGACGC | 52 | |
| | TCAATCGTCTGTCCTCTTTTGAGGACGACAAGTTTTCTTGTCGAAATGGATT | 52 | |
| | ATTTACATTGGTCCTCTTTTGAGGACGACAAGTTTTCTTGTCGCAGATTCAC | 52 | |
| | CAGTCACACGACCAGTAATAAAAGGGACAT | 30 | |
| 2 | TTACCTGAGCAAAAGAAGATGATGAAACAAACATCAAGAAAACA | 44 | 52-57 |
| | AAATTAATTAGTCCTCTTTTGAGGACGACAAGTTTTCTTGTCCATTTAACAA | 52 | |
| | TTTCATTTGAGTCCTCTTTTGAGGACGACAAGTTTTCTTGTCATTACCTTTT | 52 | |
| | TTAATGGAAAGTCCTCTTTTGAGGACGACAAGTTTTCTTGTCCAGTACATAA | 52 | |

| | | | |
|---|---|---|---|
| | ATCAATATATGTCCTCTTTTGAGGACGACAAGTTTTCTTGTCGTGAGTGAAT | 52 | |
| | AACCTTGCTTGTCCTCTTTTGAGGACGACAAGTTTTCTTGTCCTGTAAATCG | 52 | |
| | TCGCTATTAAGTCCTCTTTTGAGGACGACAAGTTTTCTTGTCTTAATTTTCC | 52 | |
| | CTTAGAATCCGTCCTCTTTTGAGGACGACAAGTTTTCTTGTCTTGAAAACAT | 52 | |
| | AGCGATAGCTGTCCTCTTTTGAGGACGACAAGTTTTCTTGTCTAGATTAAGA | 52 | |
| | CGCTGAGAAGAGTCAATAGTGAAT | 24 | |
| 3 | TGAATCTTACCAACGCTAACGAGCGTCTTTCCAGAGCCTAATTTGCCAGT | 50 | 79-85 |
| | TACAAAATAAGTCCTCTTTTGAGGACGACAAGTTTTCTTGTCACAGCCATAT | 52 | |
| | TATTTATCCCGTCCTCTTTTGAGGACGACAAGTTTTCTTGTCAATCCAAATA | 52 | |
| | AGAAACGATTGTCCTCTTTTGAGGACGACAAGTTTTCTTGTCTTTTGTTTAA | 52 | |
| | CGTCAAAAATGTCCTCTTTTGAGGACGACAAGTTTTCTTGTCGAAAATAGCA | 52 | |
| | GCCTTTACAGGTCCTCTTTTGAGGACGACAAGTTTTCTTGTCAGAGAATAAC | 52 | |
| | ATAAAAACAGGTCCTCTTTTGAGGACGACAAGTTTTCTTGTCGGAAGCGCAT | 52 | |
| | TAGACGGGAGGTCCTCTTTTGAGGACGACAAGTTTTCTTGTCAATTAACTGA | 52 | |
| | ACACCCTGAAGTCCTCTTTTGAGGACGACAAGTTTTCTTGTCCAAAGTCAGA | 52 | |
| | GGGTAATTGAGCGCTAATATCAGAGAGATAACCCACAAGAATTGAGTTAAGCC CAA | 56 | |
| 4 | TCACAAACAAATAAATCCTCATTAAAGCCAGAATGGAAAGCGCAGTCTCTGAA TTT | 56 | 106-112 |
| | ACCGTTCCAGGTCCTCTTTTGAGGACGACAAGTTTTCTTGTCTAAGCGTCAT | 52 | |
| | ACATGGCTTTGTCCTCTTTTGAGGACGACAAGTTTTCTTGTCTGATGATACA | 52 | |
| | GGAGTGTACTGTCCTCTTTTGAGGACGACAAGTTTTCTTGTCGGTAATAAGT | 52 | |
| | TTTAACGGGGGTCCTCTTTTGAGGACGACAAGTTTTCTTGTCTCAGTGCCTT | 52 | |
| | GAGTAACAGTGTCCTCTTTTGAGGACGACAAGTTTTCTTGTCGCCCGTATAA | 52 | |
| | ACAGTTAATGGTCCTCTTTTGAGGACGACAAGTTTTCTTGTCCCCCCTGCCT | 52 | |
| | ATTTCGGAACGTCCTCTTTTGAGGACGACAAGTTTTCTTGTCCTATTATTCT | 52 | |
| | GAAACATGAAGTCCTCTTTTGAGGACGACAAGTTTTCTTGTCAGTATTAAGA | 52 | |
| | GGCTGAGACTCCTCAAGAGAAGGATTAGGATTAGCGGGGTTTTGCTCAGT | 50 | |
| 5 | CAAAGTACAACGGAGATTTGTATC | 24 | 134-139 |

| | | | |
|---|---|---|---|
| | ATCGCCTGATGTCCTCTTTTGAGGACGACAAGTTTTCTTGTCAAATTGTGTC | 52 | |
| | GAAATCCGCGGTCCTCTTTTGAGGACGACAAGTTTTCTTGTCACCTGCTCCA | 52 | |
| | TGTTACTTAGGTCCTCTTTTGAGGACGACAAGTTTTCTTGTCCCGGAACGAG | 52 | |
| | GCGCAGACGGGTCCTCTTTTGAGGACGACAAGTTTTCTTGTCTCAATCATAA | 52 | |
| | GGGAACCGAAGTCCTCTTTTGAGGACGACAAGTTTTCTTGTCCTGACCAACT | 52 | |
| | TTGAAAGAGGGTCCTCTTTTGAGGACGACAAGTTTTCTTGTCACAGATGAAC | 52 | |
| | GGTGTACAGAGTCCTCTTTTGAGGACGACAAGTTTTCTTGTCCCAGGCGCAT | 52 | |
| | AGGCTGGCTGGTCCTCTTTTGAGGACGACAAGTTTTCTTGTCACCTTCATCA | 52 | |
| | AGAGTAATCTTGACAAGAACCGGATATTCATTACCCAAATCAAC | 44 | |
| 6 | ACAGTTGATTCCCAATTCTGCGAACGAGTA | 30 | 161-165 |
| | GATTTAGTTTGTCCTCTTTTGAGGACGACAAGTTTTCTTGTCGACCATTAGA | 52 | |
| | TACATTTCGCGTCCTCTTTTGAGGACGACAAGTTTTCTTGTCAAATGGTCAA | 52 | |
| | TAACCTGTTTGTCCTCTTTTGAGGACGACAAGTTTTCTTGTCAGCTATATTT | 52 | |
| | TCATTTGGGGGTCCTCTTTTGAGGACGACAAGTTTTCTTGTCCGCGAGCTGA | 52 | |
| | AAAGGTGGCAGTCCTCTTTTGAGGACGACAAGTTTTCTTGTCTCAATTCTAC | 52 | |
| | TAATAGTAGTGTCCTCTTTTGAGGACGACAAGTTTTCTTGTCAGCATTAACA | 52 | |
| | TCCAATAAATGTCCTCTTTTGAGGACGACAAGTTTTCTTGTCCATACAGGCA | 52 | |
| | AGGCAAAGAAGTCCTCTTTTGAGGACGACAAGTTTTCTTGTCTTAGCAAAAT | 52 | |

**Table S4. Sequences of the oligonucleotides to form 6x8 design (design 2) with 6 markers regions and 8 dumbbells per marker region.**

##### 3.2 Synthesis protocol for nanostructured DNA molecules

The single-stranded, circular viral DNA from M13mp18 was acquired from New England Biolabs (Catalog# N4040S). It was first linearized by hybridisation with a 39-base oligonucleotide and subsequent digestion using the BamHI and EcoRI restriction enzymes. The 39-base oligonucleotide has the following sequence

- 5’- TCTAGAGGATCCCCGGGTACCGAGCTCGAATTCGTAATC -3’

The following components were mixed first to prepare the hybridisation reaction

- 40 µL M13mp18 DNA (NEB, 250 ng/µL)
- 8 µL 10x rCutSmart buffer (NEB)

- 2 µL 39-base oligonucleotide (100 µM)
- 28 µL deionised water

The mixture was heated to 65 °C for 10 min and then linearly cooled down to 25 °C at a rate of −1 °C/min in a thermocycler. Subsequently, the following reagents were added to the reaction mixture for digestion reaction

- 2 µL BamHI-HF (100,000 units/mL)
- 2 µL EcoRI-HF (100,000 units/mL)
- 8 µL 10x rCutSmart buffer (NEB)
- 70 µL deionised water

The mixture was incubated at 37 °C for 1.5 h. The cut m13mp18 DNA was then immediately purified using the Invitrogen PureLink PCR Purification Kit (K310001) according to the manufacturer's instructions.

Following scaffold linearization, DNA nanostructures were assembled by mixing

- 20 µL cut M13mp18 DNA (final concentration ~136 nM)
- 42 µL oligonucleotide mixtures for each design (including non-replaced complementary oligo, each single oligo ~650 nM and oligo for each design, each single oligo ~1700 nM)
- 14 µL 100 mM $MgCl_2$
- 4 µL 100 mM Tris-HCl (pH 8.0), 10 mM EDTA
- 56 µL deionised water

The 136 µL mixture was heated to 80 °C for 10 min and cooled down to 70 °C, subsequently annealed using a linear cooling ramp from 70°C to 25°C over 3.5 h. As the oligonucleotides were added at 7-fold molar excess to the scaffold, the excess oligonucleotides were removed after annealing, using Amicon Ultra 50 kDa centrifugal filters. Prior to use, filters were washed multiple times with a washing buffer (10 mM Tris-HCl, 0.5 mM $MgCl_2$). Typically, one annealed sample (~136 µL) was diluted with 350 µL washing buffer and centrifuged at 9,000 g for 10 min at 4°C. The remaining volume was typically ~100 µL after centrifuge. An additional 400 µL washing buffer was then added, followed by centrifugation for another 10 min. This washing step was repeated a total of seven times.
For the final wash, 400 µL folding buffer (10 mM Tris-HCl, 2 mM $MgCl_2$, 100 mM NaCl) was added and centrifuged for 10 min. The assembled nanostructures were recovered by flushing the filter membrane on both sides with 50 µL folding buffer, followed by inverting the filter unit and centrifuging at 1,000 g for 2 min.

##### 3.3 Gel analysis of synthesized DNA nanostructures

The DNA nanostructure synthesis was characterised by agarose gel electrophoresis. All DNA samples used for the gel analysis were loaded onto a 1% agarose gel prepared in 1x TBE buffer and supplemented with 1x SYBR Safe and 2 mM $MgCl_2$. 1 kb DNA ladder (New England Biolabs, Catalog# N3232S) was used as a reference. The electrophoresis was carried out at 90 V in 1x TBE running buffer containing 2 mM $MgCl_2$ for 60 min in an ice-water bath. Gels were imaged by fluorescence imaging and UV transillumination at 15 min and 60 min, as the shorter run provides information about existing short oligonucleotides and longer run presents clear reference with the ladder sample.

To optimise the nanostructure synthesis protocol, we first characterised the product in each step, including the linearisation of plasmid via restriction enzyme cutting and nanostructure assembly by hybridisation of oligonucleotides on the linearised scaffold. For each step, samples collected before and after purification were checked to assess the effectiveness of the corresponding purification methods (Figure S10). In the 15-min run, non-purified samples after the nanostructure assembly show two bands, with the lower one representing the excess amount of oligonucleotides, and the purification effectively removed most of the oligos. In addition, in the 60-min run, the band shift observed between the samples before assembly (after plasmid linearisation) and after assembly indicates an increase in molecular size, as complementary oligonucleotides hybridise to the scaffold during annealing, nearly doubling its molecular weight.

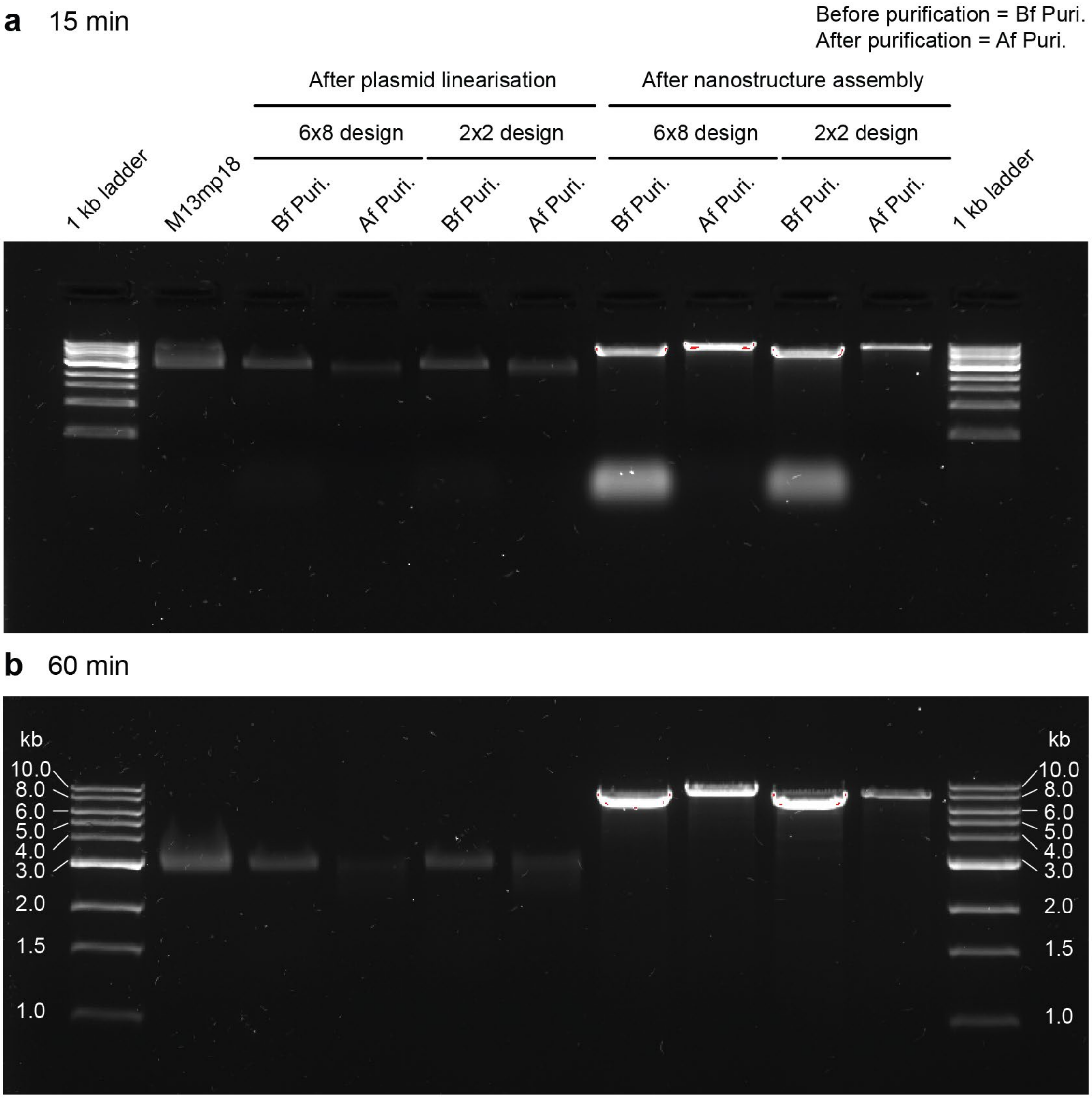

**Figure S10**. **Gel electrophoresis characterisation for quality check of each step and the purification process during DNA nanostructure synthesis. (a)** Gel running for 15 min and **(b)** 60 min in 1x TBE running buffer containing 2 mM $MgCl_2$ at 90 V. 1% agarose gel prepared in 1x TBE buffer and supplemented with 1x SYBR Safe and 2 mM $MgCl_2$ is used for the gel electrophoresis analysis.

Due to the seven-fold excess of short oligonucleotides added for dumbbell nanostructure assembly, high amounts of short DNA molecules remain in the sample and affect the translocation measurement. To reduce the oligonucleotides and avoid potential nanopore clogging, we further optimised the purification process after the assembly and increased the number of washing cycles. Gel electrophoresis analysis of the flow through (FT) after each cycle of purification shows that generally seven cycles of purification remove effectively the short oligonucleotides, which is indicated by the lower bands on the gel in 15-min run (Figure S11.a). Because the DNA concentration in the original FT was too low to produce visible bands and the concentration continued to decrease as purification cycles proceeded, the FT samples from the later purification cycles were concentrated for different lengths of time before gel analysis. This concentration process likely altered the ionic strength of the solution, which may explain the progressively slower migration of the lower bands in the subsequent FT samples (Figure S11.a). Longer run of gel electrophoresis shows the similar band shift between samples before nanostructure assembly (after plasmid linearisation) and after assembly, indicating the molecular size increase after oligo hybridisation (Figure S11.b).

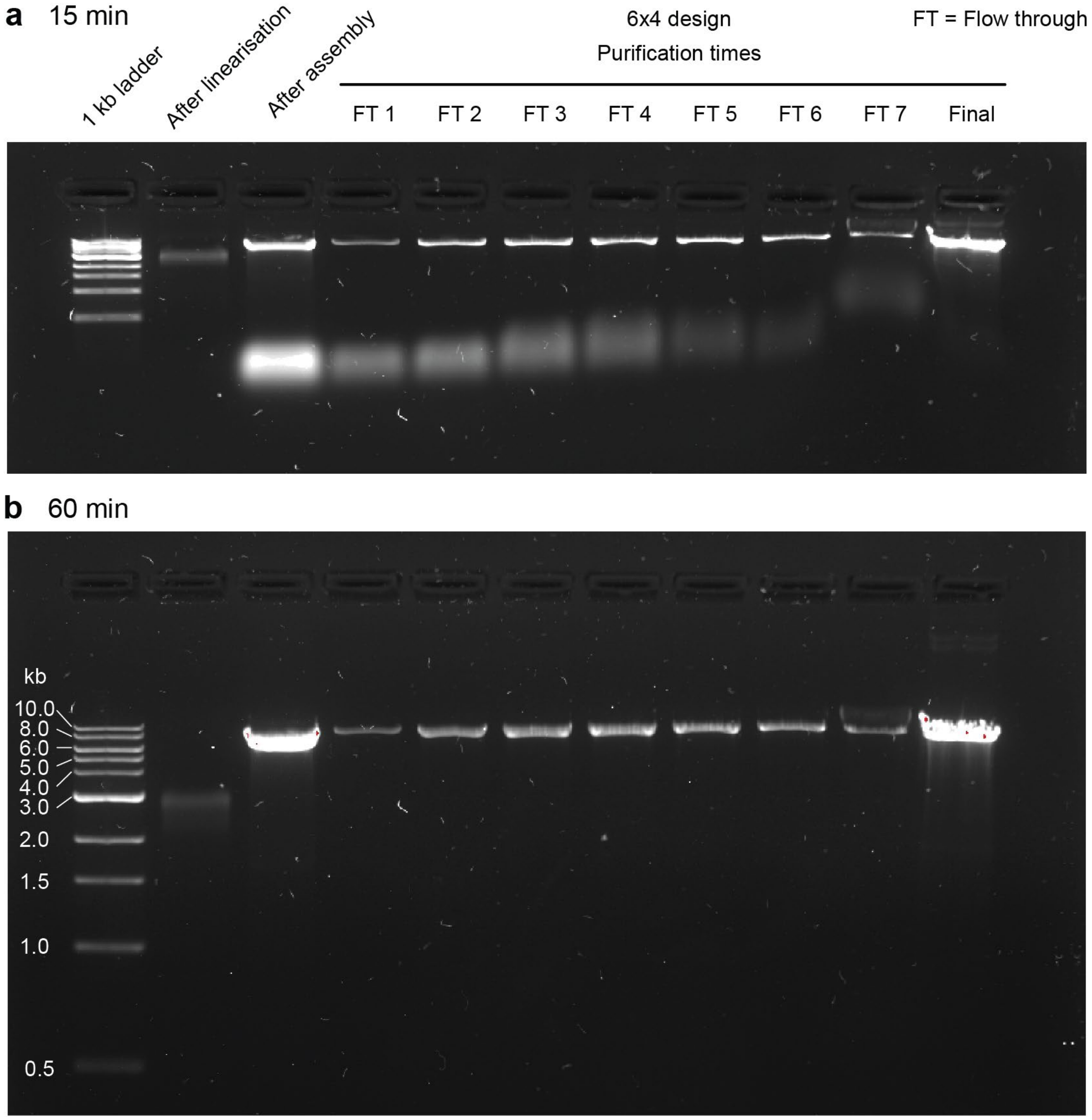

**Figure S11**. **Gel electrophoresis analysis for optimization of purification times after the nanostructure assembly process. (a)** Gel running for 15 min and **(b)** 60 min in 1x TBE running buffer at 90 V in an ice-water bath. 1% agarose gel prepared in 1x TBE buffer and supplemented with 1x SYBR Safe is used for the gel electrophoresis analysis. 'After linearisation' stands for linearised plasmid samples obtained after restriction enzyme cutting and first purification step to remove the cut fragment. 'After assembly' represents the samples obtained after hybridising oligonucleotides on the linearised plasmid scaffold while without any purification steps. The 'Final' represents samples recovered from the final wash using the storage buffer, after seven purification cycles.

##### 3.4 Stability analysis after addition of C-G pairs

The stability of dumbbell structure is important for the benchmarking test of our co-designed platform. Ideally, the hairpin structural stability should increase with more hairpin-forming base pairs and higher GC content. Table S5 shows the estimated melting temperature of two DNA sequences with one consisting of the dumbbell hairpin-forming stem in the original design and the other one used in this study, after addition of two more C-G pairs in the center of the hairpin stem. The estimation is done through the OligoAnalyzer™ tool[5] from Integrated DNA Technologies, Inc. (IDT) using approximate condition of final sample in storage buffer with around 10 nM concentration in 100 mM NaCl, 2 mM $MgCl_2$ solution. After adding the 2 C-G pairs in the center, the new design with 12 bp shows much higher melting temperature compared with the original design with 10 bp. It is worth marking that the melting temperature estimation using only the pairing segment is different from that of a self-pairing hairpin, and the result therefore only reflects the higher stability of the new design after the addition of C-G pairs.

| Design | DNA Sequence | Basepair length | Melting Temperature |
|---|---|---|---|
| Without adding C-G pairs | 5'- GAGGAACAAG -3'<br>5'- CTTGTTCCTC -3' | 10 | 25.1 ℃ |
| After adding C-G pairs | 5'- GAGGA**CG**ACAAG -3'<br>5'- CTTGT**CG**TCCTC -3' | 12 | 38.8 ℃ |

**Table S5. Estimation of melting temperature for two designs of DNA sequences using the self-pairing oligonucleotide sequence of the dumbbell stem.** The melting temperature is estimated using the OligoAnalyzer™ tool (IDT, Inc.) based on the pairs formed by the listed sequence with its complement sequence, with settings of 10 nM oligonucleotide concentration and 100 mM NaCl salt solution with 2 mM $MgCl_2$, similar to the sample storage buffer.

The thermal stability of the self-pairing hairpin is then estimated using the OligoAnalyzer™ tool[5] under the same setting but with input of single oligonucleotide sequences that form the complete dumbbell unit. Figure S12 shows that addition of 2 C-G pairs in the center of the dumbbell stem increases the thermal melting temperature by around 5 ℃. In addition, the C-G pair in the center of dumbbell structure provides a stronger binding compared with the A-T pairs in the original design, when under the strain exerted by the extended 5' and 3' oligonucleotide backbone outside the dumbbell region (i.e., 5'-TTG and 3'-TCC). This new design therefore presents higher stability for each single dumbbell unit.

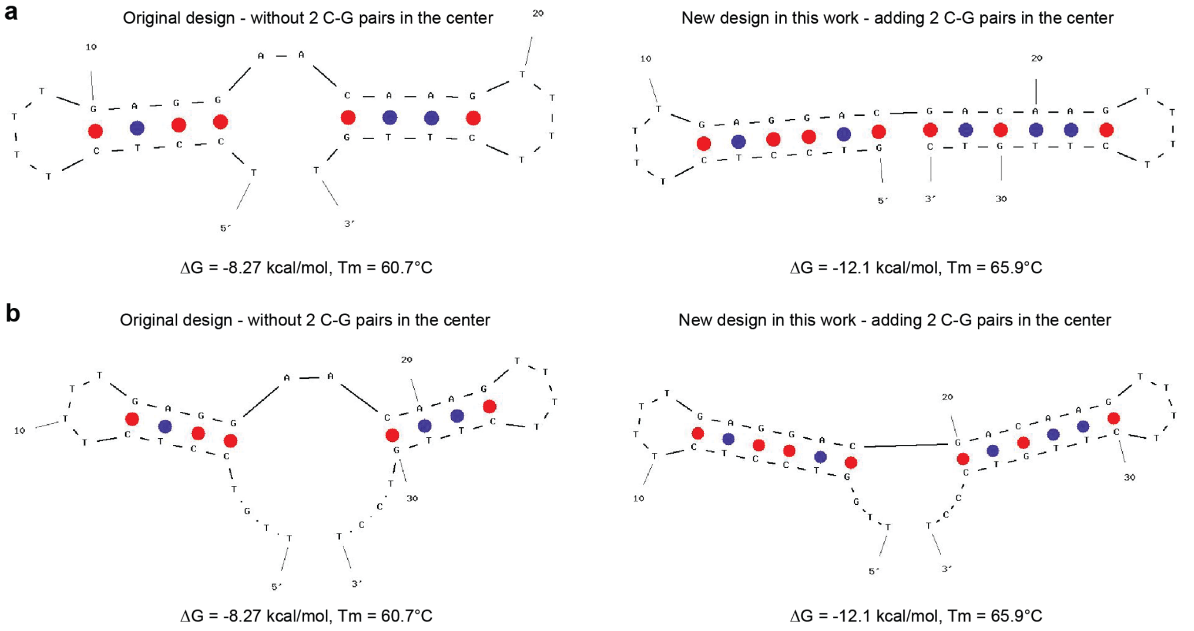

**Figure S12. Simulated structure and melting temperature of the two designs, including one original design without adding C-G pairs (left column) and new design in this study after adding C-G pairs (right column).** The simulation was performed on the different segments of the oligonucleotides that form the dumbbell unit, with **(a)** containing only dumbbell part, and **(b)** containing 3 more nucleotides extension on each end of the oligonucleotide (5'-TTG and 3'-TCC) as a reference to approximate the real case. The sequence of two extended tails in (b) is the same as that of oligonucleotide used for synthesis. Numbers such as 10, 20 and 30 indicate the nucleotide position in the oligo sequence.

To verify the formation of dumbbell unit in the new design, we used NUPACK[2,3] to analyze the probable structures by inputting both the dumbbell unit-containing oligonucleotide sequence and the corresponding sequence of the single-strand DNA backbone, simulating the final dsDNA molecule that carries the dumbbell structures in a final concentration of 10 µM. The result in Figure S13 shows the proxy structures of both single dumbbell unit and single complete marker region in the 6x4 and 6x8 designs that have minimum free energy, with all pairs presenting close to 100% equilibrium probability. This therefore provides theoretical support for the structure of synthesised molecules after the modification on the redesign of the dumbbell unit.

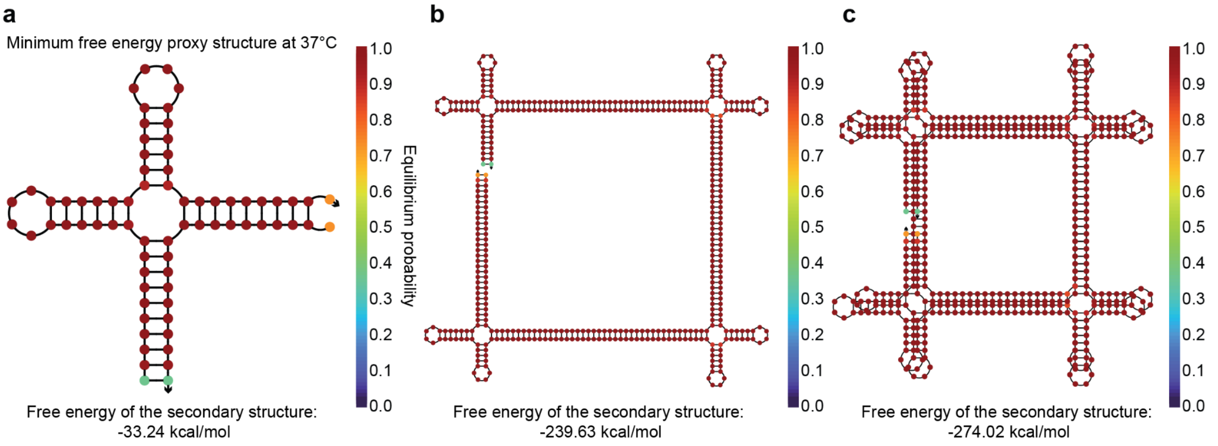

**Figure S13.** Simulated minimum free energy (MFE) proxy structure[2] for **(a)** a single dumbbell unit, **(b)** a single marker region for 6x4 design with 4 dumbbell units, and **(c)** a single marker region for 6x8 design with 8 dumbbell units, after adding the 2 C-G pairs in new design.

#### Section 4. Nanopore measurement and data analysis

As nanopore experiments record digital signals after the analog-to-digital conversion, ionic current and time series are obtained as raw data, which is processed in a few steps until the statistical information is obtained. The acquisition and analysis workflow is described as follows.

**Signal Acquisition and Pre-processing:** The Ionic current recordings are acquired at a sampling rate of 40 MHz (25 ns per sample). Two signal representations are maintained in parallel throughout the analysis. A 2 MHz Bessel low-pass filter is applied when extracting the dumbbell-associated DNA translocation events from the raw trace through Poriscope[6] and the low-pass filtered reference signal (referred to as "2 MHz LP-filtered reference"), serves as a noise-reduced trace for event segmentation and marker detection. Separately, applying the deep-learning (DL) U-Net denoiser to the raw signal produces a reconstructed current trace (referred to as "DL reconstruction") that preserves sub-microsecond features while suppressing broadband noise. Both representations share the same time axis sampled at 40 MHz.

**Baseline Noise Estimation and Event Body Detection:** The pre-event baseline for each event is extracted from the 2 MHz LP-filtered reference, using the segments preceding the translocation onset. The noise level is quantified using the median absolute deviation (MAD) estimator

$$\hat{\sigma} = 1.4826 \times median(|x - median(x)|)$$

where $x$ denotes the baseline current samples, and the constant 1.4826 is a rescaling factor that makes $\hat{\sigma}$ an asymptotically consistent estimator of the standard deviation under Gaussian noise while remaining robust to spike-like outliers. This robust noise estimate is used as the detection sensitivity parameter in all subsequent threshold calculations. Instead of using standard deviation directly, we adopted the MAD method as the real baseline segments could deviate from Gaussian due to either capacitive transients at the start of recording or after voltage steps, or the occasional sharp artifacts, which therefore makes standard deviation sensitive to outliers. Boundaries of each translocation event are identified as the event start and event end using the open-pore baseline level and five-fold threshold. The open-pore baseline level is estimated as the median of the first 200 samples of the 2 MHz LP-filtered reference.

**Event Segmentation (Stage 1, Coarse Detection):** To detect individual DNA translocation events, an additional 1 MHz fourth-order Butterworth low-pass filter was applied to the 2 MHz LP-filtered reference using zero-phase forward–backward filtering. The mean of the filtered signal over the entire event window was computed as the backbone current level. A detection threshold was defined as:

$$\theta = mean_{backbone} - 5 \times \hat{\sigma}$$

Contiguous segments that fall below the threshold are then identified as candidate marker windows, with defined start of end time for each window.

**Dumbbell Peak Detection within Marker Windows (Stage 2, Fine Detection):** Peak detection is performed on the DL reconstruction signal within each detected marker window, exploiting the higher temporal resolution and well-preserved transient details in the signal. Peak-finding strategy is applied based on find_peaks function in Scipy package, using a prominence threshold of $1.0 \times \hat{\sigma}$ to extract local peak feature on the denoised signal. The 2-stage detection process is shown in Figure.S14.

**Peak Accumulation for Statistics:** Detected peak times (in μs) are stored together with their associated spacings and two measurements are computed. The first computation is the intra-marker peak-to-peak interval, which is defined as the time interval between consecutive peaks within a dumbbell pair (for example, $d_{01}$ and $d_{23}$ for the 2x2 design). Intra-marker translocation velocity is then computed using the intra-marker time and the known physical spacing of the dumbbell peaks in base pairs (bp) as determined by the DNA design. The intra-marker interval and velocity distribution from all the events are plotted in log-scale histogram and the mean and standard deviation values are extracted from the fitting and reported in Figure.3,4 in the Main text.

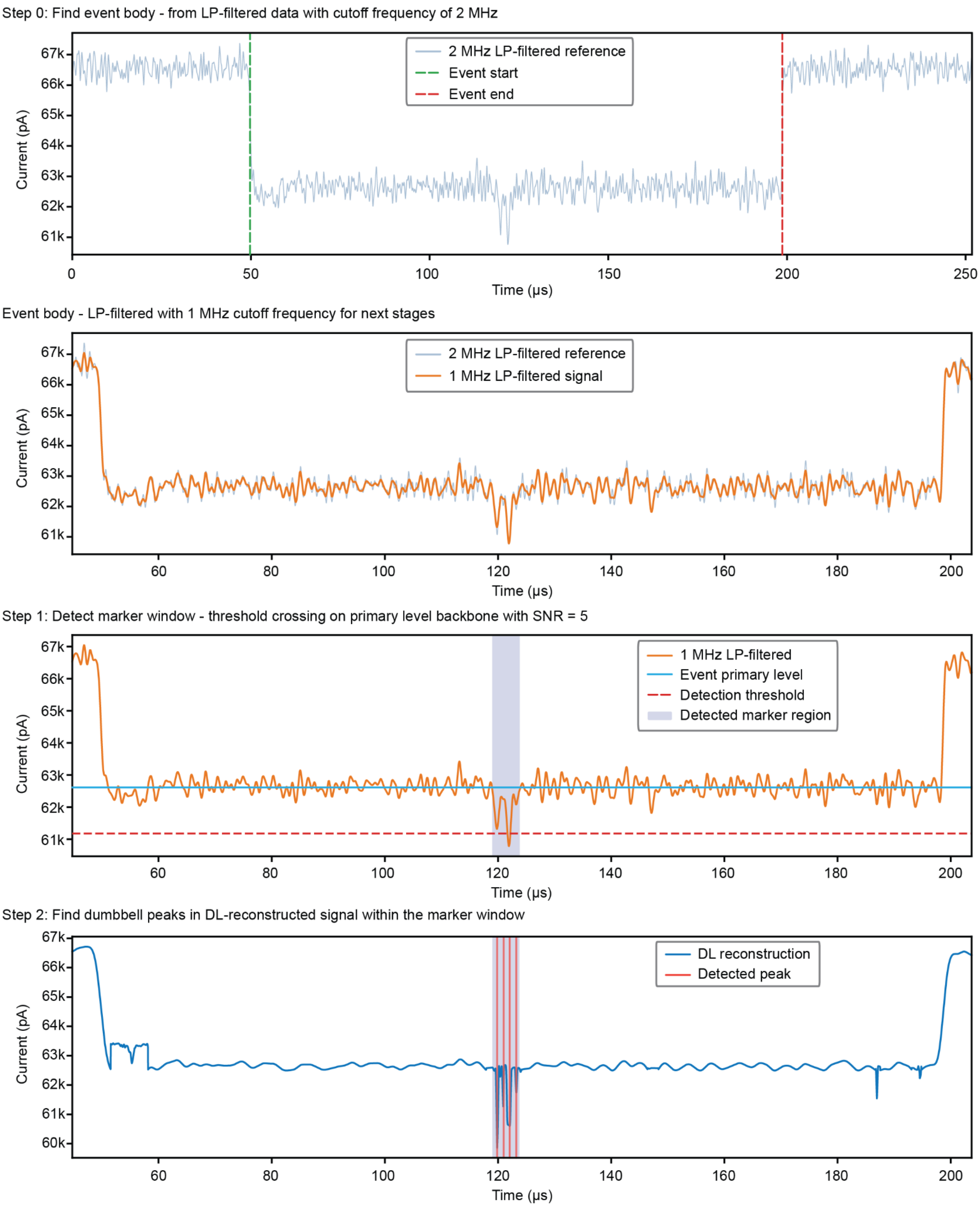

**Figure S14. Stepwise illustration of peak-finding workflow.** The event body is defined as the segment where current blockage happens owing to molecule translocation. We first detect the event body using a threshold-crossing method based on the LP-filtered data to ensure sufficient signal-to-noise ratio (SNR). The event body is then analysed in two steps to first detect the marker region using a threshold-crossing method on 1 MHz LP-filtered trace (step 1), and the dumbbell peaks are then located inside the marker region using the peak-finding algorithm on DL-reconstructed signal (Step 2).

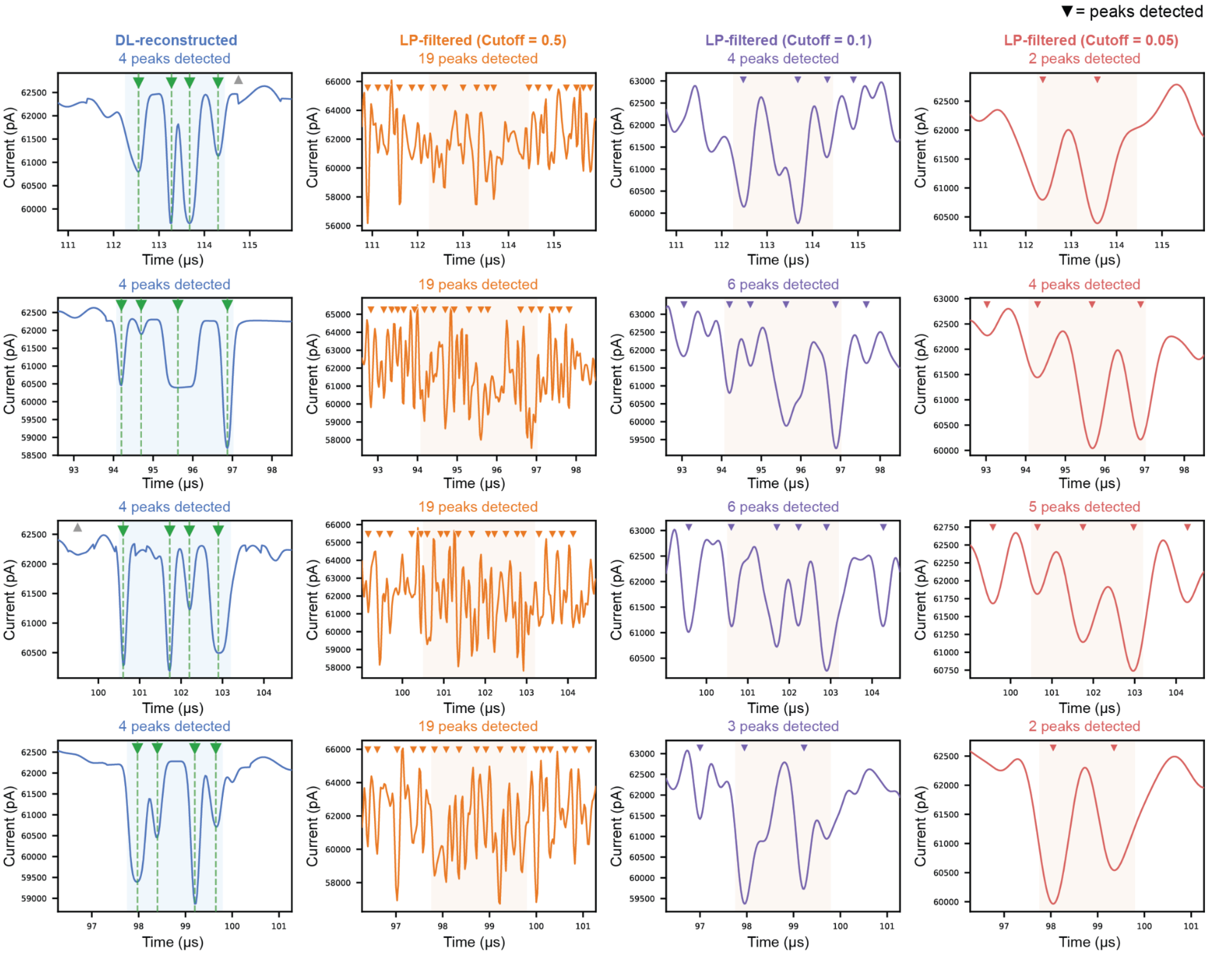

**Figure S15. Comparisons of DL reconstruction and LP-filtering in terms of peak calling efficacy**. The marker region of 4 example events measured from 2x2 design are shown. A same peak-finding algorithm is applied on all input signals, including DL-reconstructed signals and LP-filtered signals with different cutoff levels. Detected peaks on various input signals are indicated with triangles and dashed lines. Compared with LP-filtered signals where the peak quality degrades and peak detection results vary with cutoff frequency, reconstructed signals using the neural network display clearly resolved peaks with sharper temporal boundaries and higher SNR.

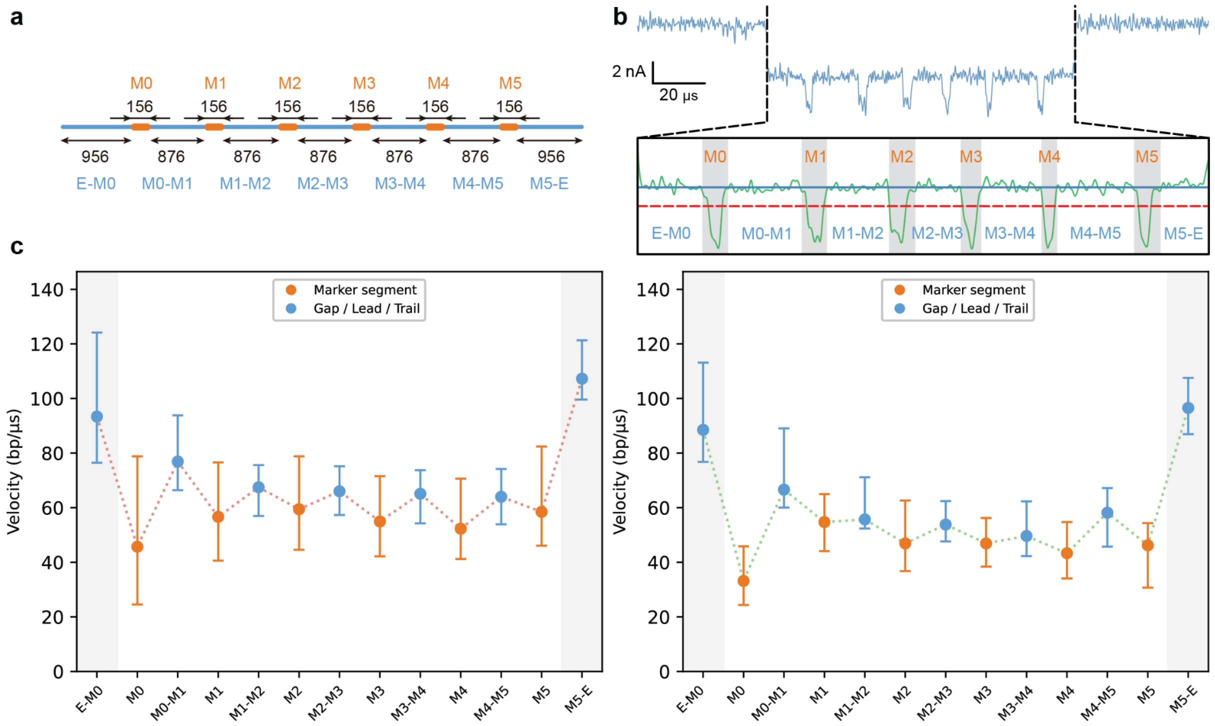

**Figure S16. Individual molecule translocation velocity profiles for DNA constructs of 6x4 and 6x8 designs. (a)** Schematic representation of a DNA molecule carrying 6x8 design, with indication of marker segments and inter-marker segments (gaps, lead or trail). Contour distances in basepairs (bp) are estimated according to the sequence of designed construct. **(b)** Demonstration of the threshold-crossing algorithm used to extract segmental duration from the 2 MHz LP-filtered current signal. **(c)** The translocation velocity profiles obtained from 6x4 design (left) and 6x8 design (right). Error bars correspond to the standard deviation of velocity among all events used for statistical analysis (n = 189 for 6x4 design, n = 29 for 6x8 design).

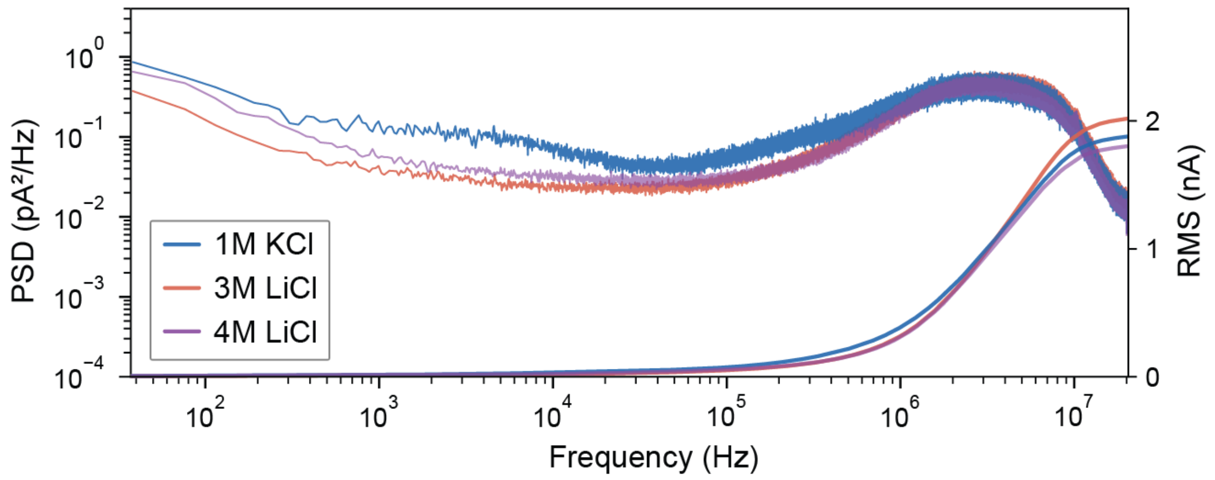

**Figure S17. Noise comparison under different measurement conditions.** Three buffer solutions used in this work are compared, including 1 M KCl (10 mM Tris-HCl and 1 mM EDTA, pH 8.0), 3 M LiCl (10 mM Tris-HCl and 1 mM EDTA, pH 8.0) and 4 M LiCl (10 mM Tris-HCl and 1 mM EDTA, pH 9.0). Ionic currents are collected using separate low-noise chips with nanopore diameters of 8.0 nm, 8.6 nm and 6.5 nm for 1 M KCl, 3 M LiCl and 4 M LiCl, respectively. All current signals are measured under 500 mV voltage bias using the 10 MHz amplifier from Elements. The 1 M KCl shows slightly higher noise compared with the LiCl and noise generally reduces with increasing salt concentration, in accordance with the previous report[7,8].

| | 2x2 design | 6x4 design | 6x8 design |
|---|---|---|---|
| CH1 | 101 ± 34 | 103 ± 20 | 92 ± 37 |
| CH2 | 84 ± 34 | 69 ± 13 | 108 ± 44 |

**Table S6. Peak detection rate (%) for three different molecule designs in each individual channel.** The detection rate is calculated by dividing the number of detected peaks in each event over the total peak numbers from the design of DNA constructs.

| | 2x2 design | 6x4 design | 6x8 design |
|---|---|---|---|
| Marker recall rate | 100 ± 0 | 100 ± 0 | 98 ± 4 |
| Peak recall rate | 77 ± 24 | 66 ± 13 | 78 ± 16 |

**Table S7. Cross-channel marker and dumbbell-peak recall rates (%) for all three different molecule constructs.** Both recall rates are defined as whether corresponding marker or individual dumbbell-peak is detected in CH2 when the marker or peak is detected in CH1.

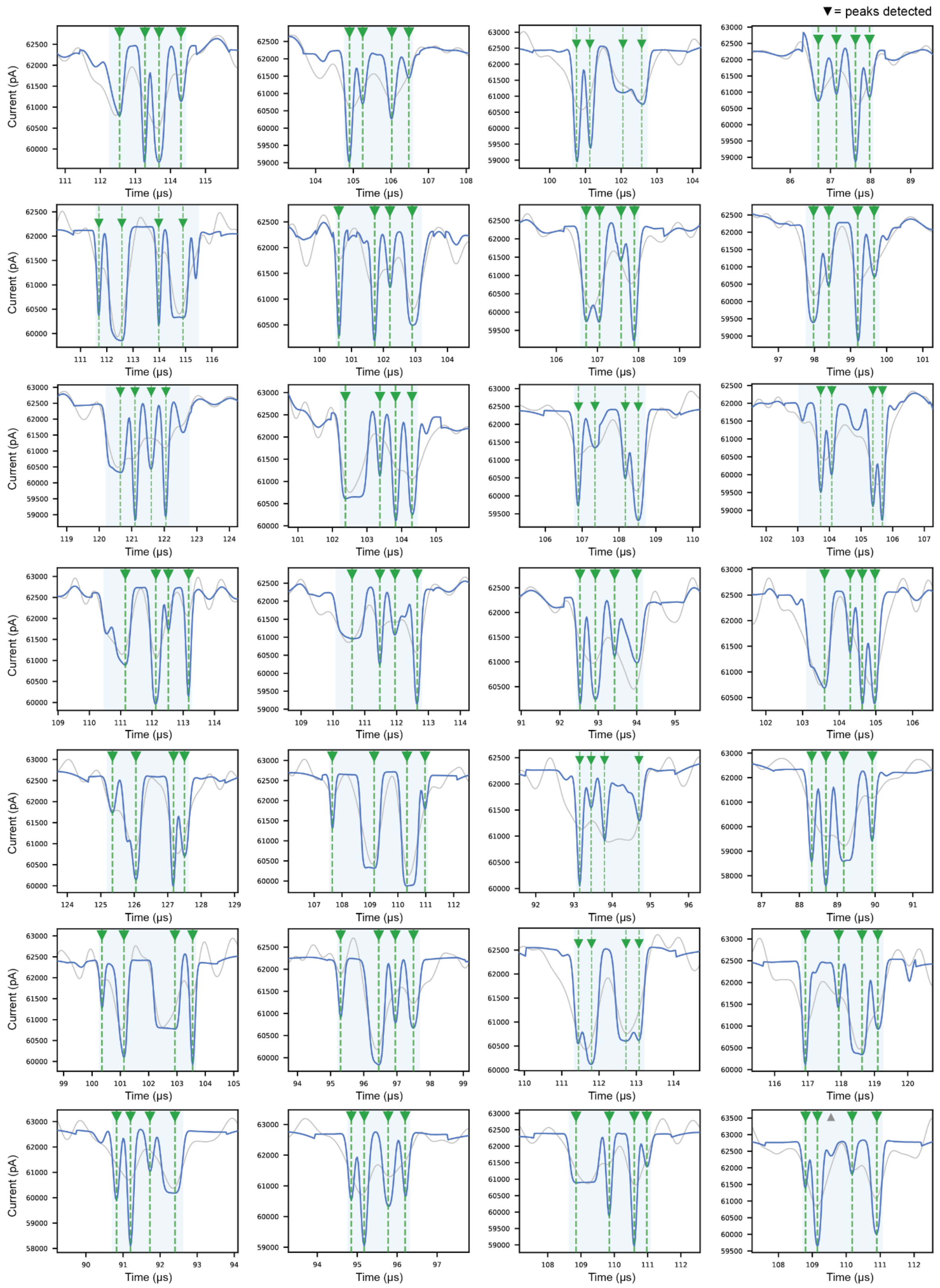

**Figure S18. Example events from 2x2 design after DL-reconstruction.** Both DL-reconstructed signals (blue) and digitally LP-filtered signals (grey, cutoff frequency at 2 MHz) of the marker region are shown. The marker region contains 4 dumbbells. Detected marker regions are highlighted in light green shade, and peaks detected by the peak-finding algorithm are indicated by green triangles and dashed lines. Using the 2 MHz digitally LP-filtered signal as a reference, the DL-reconstructed results exhibit clearly resolved peaks with well-defined temporal boundaries and improved SNR.

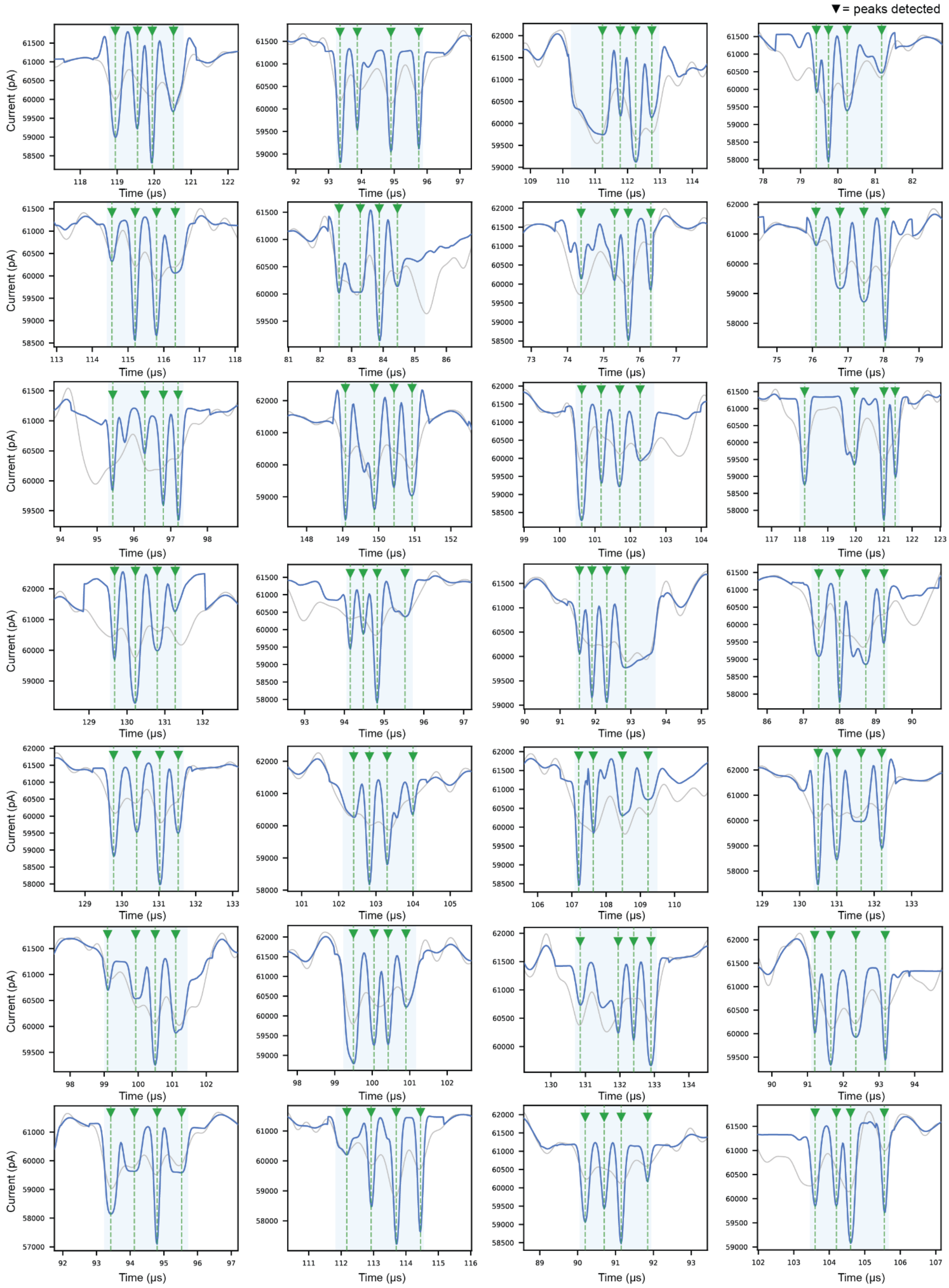

**Figure S19. Example events from 6x4 design after DL-reconstruction.** Both DL-reconstructed signals (blue) and digitally LP-filtered signals (grey, cutoff frequency at 2 MHz) of individual marker regions are shown. Each marker contains 4 dumbbells by design. Detected marker regions are highlighted in light green shade, and peaks detected by the peak-finding algorithm are indicated by green triangles and dashed lines.

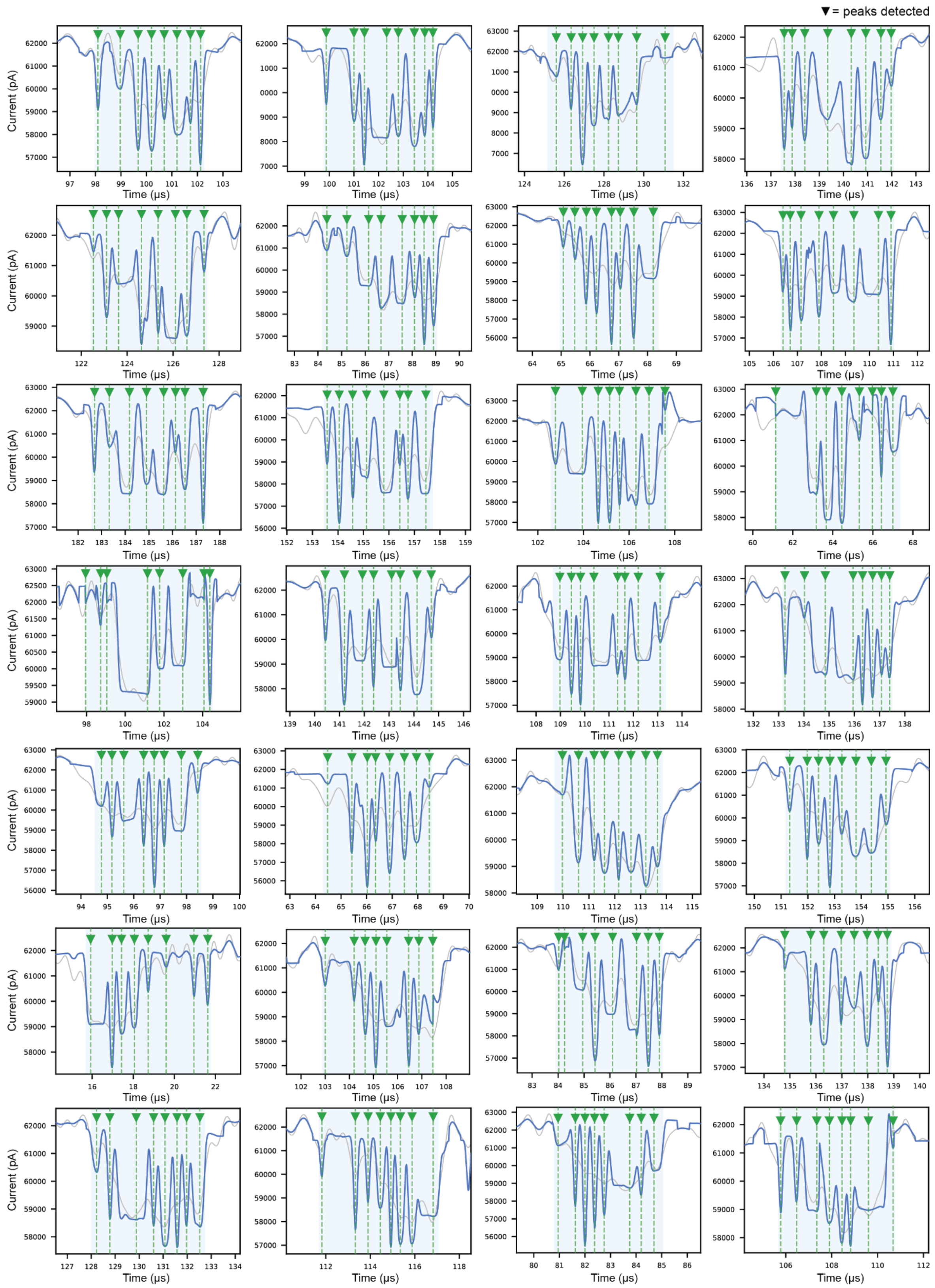

**Figure S20. Example events from 6x8 design after DL-reconstruction.** Both DL-reconstructed signals (blue) and digitally LP-filtered signals (grey, cutoff frequency at 2 MHz) of individual marker regions are shown. Each marker contains 8 dumbbells by design.

Detected marker regions are highlighted in light green shade, and peaks detected by the peak-finding algorithm are indicated by green triangles and dashed lines.

#### References

1. Chen, K. *et al.* Super-Resolution Detection of DNA Nanostructures Using a Nanopore. *Advanced Materials* **35**, 2207434 (2023).
2. Zadeh, J. N. *et al.* NUPACK: Analysis and design of nucleic acid systems. *Journal of Computational Chemistry* **32**, 170–173 (2011).
3. Fornace, M. E. *et al.* NUPACK: Analysis and Design of Nucleic Acid Structures, Devices, and Systems. https://doi.org/10.26434/chemrxiv-2022-xv98l-v2 (2025) doi:10.26434/chemrxiv-2022-xv98l-v2.
4. Chen, K. *et al.* Dynamics of driven polymer transport through a nanopore | Nature Physics. *Nature Physics* **17**, 1043–1049 (2021).
5. Owczarzy, R. *et al.* IDT SciTools: a suite for analysis and design of nucleic acid oligomers. *Nucleic Acids Res* **36**, W163–W169 (2008).
6. González González, A. C., Kerrouri, N., Wadhwa, D., Tabard-Cossa, V. & Briggs, K. Poriscope: A Configurable Pipeline for Nanopore Data Analysis. *Journal of Open Research Software* **14**, (2026).
7. Smeets, R. M. M., Keyser, U. F., Dekker, N. H. & Dekker, C. Noise in solid-state nanopores. *PNAS* **105**, 417–421 (2008).
8. Saharia, J., Bandara, Y. M. N. D. Y., Karawdeniya, B. I., Alexandrakis, G. & Kim, M. J. Assessment of 1/f noise associated with nanopores fabricated through chemically tuned controlled dielectric breakdown. *ELECTROPHORESIS* **42**, 899–909 (2021).
